# Xlink Mapping and AnalySis (XMAS) - Smooth Integrative Modeling in ChimeraX

**DOI:** 10.1101/2022.04.21.489026

**Authors:** Ilse M. Lagerwaard, Pascal Albanese, Andris Jankevics, Richard A. Scheltema

**Affiliations:** Biomolecular Mass Spectrometry and Proteomics, Bijvoet Center for Biomolecular Research and Utrecht Institute for Pharmaceutical Sciences, Utrecht University, Padualaan 8, 3584 CH Utrecht, The Netherlands; Netherlands Proteomics Centre, Padualaan 8, 3584 CH Utrecht, The Netherlands

**Keywords:** ChimeraX, XL-MS, XlinkX/PD, PhoX, DSSO, DSS, fibrin

## Abstract

Crosslinking mass spectrometry (XL-MS) adds tremendous value to structural biology investigations. Its main strength lies in uncovering structural information in the form of distance constraints between neighboring amino acids of proteins, protein regions and complex samples, which are difficult to assess by other analytical techniques. However, although several approaches have been proposed, interpreting XL-MS data in a structural context has been cumbersome. ChimeraX has gained momentum as a flexible and widely used software package for the visualization of structural data, but is currently lacking functionalities for integration of experimental XL-MS data. Here, we introduce XMAS, a bundle that allows users to load results from several XL-MS search engines directly into ChimeraX and map the information onto protein structures. Besides automatically locating distance constraints on protein structures, XMAS offers the possibility to work with replicate experiments and/or different crosslinkers, and filter this data based on the number of replicates for which a given distance constraint was detected, thereby increasing the data quality. Additionally, we introduce the concept of self-links, which allows easy modeling of homo-dimeric interactions. Its core functionality is extended by the implementation of seamless connections to the HADDOCK suite to streamline otherwise time-consuming tasks in structural modeling pipelines. We demonstrate these key elements of the XMAS bundle by modeling crosslinking data obtained from human fibrin clots. The software is freely available from the ChimeraX toolshed, with an extensive user manual and example datasets.

---

Proteins are large, macro-molecular machines responsible for highly complex cellular behavior such as sensing the external environment, reproduction, defense, and many others. They do not always act alone, but also cooperate in larger assemblies that enable more complex behavior than the sum of the individual parts would suggest (1). It has been estimated that 80% of the proteins engage in these interactions, leading to estimations of 130,000 to 650,000 unique protein to protein interactions (2, 3). The study of protein structure and the protein complexes they form is an important field, requiring solid analytical approaches, that aids our understanding of for example complex diseases driven by failures in the protein machinery. A powerful technique to analyze proteins is shotgun mass spectrometry, where proteins are digested into peptides before separating the peptide mixture by liquid chromatography and measuring the eluting peptides through mass spectrometry (4). The technique has proved instrumental in uncovering relative quantities of proteins between different cellular states (5, 6, 7), location of post-translational modifications (6), melt curves of proteins (7, 8), location in cellular compartments (8, 9, 10), and many other details of the life of proteins inside and outside cells. To make the technique capable of capturing protein structural information, XL-MS was developed (9, 10, 11). Here, agile reagents that are capable of forming a covalent bond between amino acids in close proximity are supplemented to the protein mixture. Typically, NHS ester chemistry is utilized to capture lysine-lysine pairs but other chemistries have also been introduced (11, 12). After digestion, the peptide mixture contains three distinct products: normal peptides (unaffected by the used reagent), mono-links (a single peptide where one amino acid is covalently linked by one reactive head on the used reagent, while the other quenched on for example water), and cross-linked peptides (two peptides covalently bound by the reagent). The term crosslink is then defined as two connected amino acids, independently from the presence of missed cleavages or modifications in the containing peptides (13). The experimentally derived crosslinks are of interest for structural studies, as from the peptide identities distance constraints can be derived delineating the structure of individual proteins and the interaction interface(s) between multiple proteins. In recent years, the technique has been made generally applicable by many developments that specifically solved issues with low stoichiometry of the desired products and complexities in data analysis (14–17). Aiding its adoption was the advent of the cryo-electron microscopy (EM) era in structural biology where researchers found XL-MS to be a perfect companion (17), enabling the interpretation of Cryo-EM/ET densities where the proteins cannot be constructed *ab initio* or the identity of interacting subunits is difficult to recover (18, 19).

More recently, XL-MS has shown great potential for leveraging the spectacular advances in *in-silico* protein prediction solutions like AlphaFold2 (20) and RoseTTAFold (21) and even the recently introduced ability to use AlphaFold2 to gain insights into protein complexes (22, 23). Using the distance constraints, the produced models can be validated for correctness and utilizing structural modeling approaches like HADDOCK (24) aids in the building of credible structural models of protein complexes. Visualization of distance constraints and their use inside the 3-dimensional protein structures is critical in this respect and has received considerable attention in the past. For example, tools were developed to visualize the distance constraints in plots showing the primary protein structure to gain insights into hotspots of crosslinking, type of folding and many other details (25–29). These approaches have enabled scientists to make use of the structural information derived from XL-MS experiments, but at a price. Data (re-)formatting and additional scripting have been unavoidable for many years leading to a steep learning curve and, importantly, potentially leading to mistakes in data interpretation.

Here, we introduce XMAS, a plugin that is fully integrated in ChimeraX (30) that allows data from XL-MS experiments – generated from multiple search engines – to be directly loaded into the environment without the need for data formatting. Additionally, the plugin automatically localizes the crosslinks inside the protein sequence / structure even in those cases where the sequences do not perfectly match (*e.g*. when only a part of the protein is structurally resolved). We also introduce the concept of self-links that assist – even in existing XL-MS datasets as no additional experimental setup is required – to build protein complex models for homomers. Furthermore, automatic searches for the shortest distance for cases where multiple locations are possible (e.g. for homomers), plotting of distances, and overlap between replicate experiments are provided. Lastly, to enable full use of the distance constraints in a structural modeling context, XMAS automates the connection to the HADDOCK structural modeling environment (24, 31). This takes away a large amount of data (re-)formatting steps and eases their use for nonexpert users. As supplementary material, we provide an extensive manual and practice data (see section Supplementary Material).

## Materials and methods

### Implementation

XMAS was implemented as a ChimeraX bundle written in Python 3.9. It was packaged as a Python wheel according to the ChimeraX developer guide for Building and Distributing Bundles, which can be found at https://www.cgl.ucsf.edu/chimerax/docs/devel/writing_bundles.html. Of the third-party Python packages pre-included in the ChimeraX environment, XMAS uses PyQt5 (for the graphical user interface (GUI)), NumPy (for fast processing of large datasets), and Matplotlib (for generating plots). Additionally, Pandas (for handling tabular data format files), openpyxl (for importing Excel files), matplotlib-venn (for plotting Venn diagrams), seaborn (for plotting box and strip plots), and QtRangeSlider (to create slider widgets for defining a range of values) are employed. These external libraries are automatically installed in ChimeraX upon installation of XMAS when necessary.

XMAS can be downloaded and installed easily via the ChimeraX toolshed at https://cxtoolshed.rbvi.ucsf.edu/. Alternatively, instructions for manual installation are provided in the ‘INSTALL.TXT’ file included in the install package. To ensure support of all XMAS functionalities, ChimeraX version 1.3 or higher needs to be installed. The bundle source code, including a user manual, is freely available at https://github.com/ScheltemaLab under an Apache License 2.0 software license.

### Crosslink Datasets

Commercial plasminogen-depleted fibrinogen, purified from human plasma (Merck), was dissolved in ice-cold crosslinking buffer at a concentration of 10 μM (1mg/mL, 100 μg per 1.5mL vial). The crosslinking buffer contained 50 mM Hepes (pH 7.4), 120 mM NaCl, and 2 mM CaCl_2_ (all Sigma-Aldrich). Fibrinogen clotting was activated by adding 1 U/mL human thrombin (Sigma-Aldrich). Samples were incubated at 37°C for 30 min in an Eppendorf Thermomixer C orbital shaker to allow further clotting. Two sets of three independently clotting reactions on 100 μg of fibrinogen each were crosslinked at room temperature for 30 min with two different crosslinking reagents: 2 mM (3,5-bis(((2,5-dioxopyrrolidin-1-yl)oxy) carbonyl)phenyl)phosphonic acid (PhoX) or 1 mM disuccinimidyl suberate (DSS) (Thermo Scientific). The crosslinking reaction was subsequently quenched with 50 mM Tris (Roche). Next, pre-digestion was performed by incubation with Lys-C (Sigma-Aldrich) at 37°C for 30 min, at a Lys-C/protein ratio of 1:70 (wt/wt). Proteins were denatured with 4 M urea (Merck) at 54°C for 1 h, prior to sonication in a Bioruptor bath sonicator (Diagenode) for 15 cycles of 30 s ON and 30 s OFF. Thereafter, samples were reduced with 8 mM DTT (Sigma-Aldrich) at 54°C for 30 min in an Eppendorf Thermomixer C at 800 RPM, followed by alkylation with 200 mM IAA (GE Healthcare) at room temperature for 1 h while protected from light exposure. Samples were then diluted with 50 mM Tris at a sample/Tris ratio of 1:1.1 (vol/vol) to reduce the urea concentration to <2M. Trypsin and Lys-C ( Sigma-Aldrich) were added at a protease/protein ratio of 1:25 and 1:70 (wt/wt), respectively, for protein digestion overnight at 37°C (16h). Prior to peptide cleanup, samples were acidified to pH < 2 with 0.5% trifluoroacetic acid (TFA) (Honeywell) and desalted using an Oasis HLB μElution 30 μm plate (Waters). After washing the plate columns, twice with 150 μL of acetonitrile (ACN) (Biosolve) and twice with 150 μL of 0.1% TFA, acidified samples were loaded. Columns were then washed twice with 150 μl 0.1% TFA again, and peptides were eluted with a 50% ACN/0.1% TFA (thrice 35 μl). The obtained eluates were evaporated by vacuum centrifugation in a Savant SpeedVac Vacuum Concentrator (Thermo Scientific). For subsequent fractionation of crosslinked peptides, two different strategies were adopted, depending on the chemical crosslinker. For PhoX crosslinked samples, the previously optimized enrichment protocol leveraging the IMAC enrichable phosphonate group was used (14). Briefly, crosslinked peptides were enriched with Fe(III)-NTA-5 μL using the AssayMAP Bravo Platform (Agilent Technologies). Cartridges were primed with 200 μL of 0.1% TFA in ACN and equilibrated with 200 μL of 80% ACN/0.1% TFA. Samples were dissolved in 200 μL of 80% ACN/0.1% TFA and loaded onto the cartridge. The columns were washed with 250 μL and the crosslinked peptides were eluted with 35 μL of 10% ammonia directly into 35 μL of 10% formic acid (FA) (98-100% purity; Merck). Samples crosslinked with DSS were enriched by strong cation exchange (SCX) chromatography using the same AssayMAP Bravo Platform with a protocol specifically optimized for the purpose of micro-fractionation of crosslinked peptides pairs, as follows: dried samples were resuspended in 100 μL 20% ACN/0.1% TFA, using SCX cartridges in an AssayMAP Agilent Bravo robot (both Agilent Technologies). Fractions were retrieved by five consecutive elutions of 25 μL each with buffers containing increasing concentrations (50, 250, 450, 600, and 750 mM) of ammonium acetate (diluted from a 7.5 M stock; Sigma-Aldrich), in addition to 20% ACN, and TFA to acidify buffers to pH < 2. Next, the 250-, 450-, 600-, and 750-mM ammonium acetate SCX fractions were desalted on an Oasis HLB μElution 30 μm plate as described previously in this section. All fractions were evaporated again in a Savant SpeedVac Vacuum Concentrator and stored at −20°C prior to MS analysis.

For MS acquisition, samples were resuspended in 2% formic acid and peptides were separated using an Ultimate 3000 LC system (Thermo Scientific) coupled online to an Orbitrap Fusion Lumos Tribrid mass spectrometer (Thermo Scientific). Firstly, peptides were trapped in reverse-phase solvent A (0.1% FA), by applying a 5-μL/min column flow rate for 5 min on a 2-cm double-frit trap column (100-μm inner diameter) packed in-house with Reprosil-Pur C18-AQ 3 μm beads (Dr. Maisch). Secondly, peptide separation was achieved by a 45-min gradient elution with 9-42% solvent B (80% ACN/0.1% FA), applying a 300-nL/min column flow rate on a 50-cm single-frit analytical column (75-μm inner diameter) packed in-house with InfinityLab Poroshell 120 EC-C18 2.7 μm beads (Agilent Technologies).

Survey MS1 Orbitrap scans were acquired at 60,000 resolution, with an automatic gain control (AGC) target of 1e6 ions, maximum injection time (IT) of 50 ms, and scan range of 375-1500 *m/z*. Precursors with a 3-8 charge state were selected for fragmentation using a stepped HCD collision energy mode (normalized energies of 24-27-30% for DSS and 19-27-35% for PhoX). Dynamic exclusion properties were set to sort by highest charge state. Full MS2 Orbitrap scans were acquired at 30,000 resolution, with an AGC target of 8e5 ions and dynamic maximum IT. Spectra were analyzed using Thermo Proteome Discoverer 2.4.1.15 software with incorporated XlinkX nodes, using DSS (32) or PhoX (14) specific settings. Crosslinks with a false discovery rate at 1% and score >40 were kept for further analysis. Unless otherwise specified, only crosslinks present in at least 3 out of 3 replicates were used for structural modeling(32).

### Structural Modeling

As a proof of concept for testing XMAS to uncover novel structural insights, we used as a starting model the laterally aggregating fibrinogen protofibrils from Klykov *et al*. (33) that was based on the available X-ray crystal structure (PDB ID: 3ghg). All additionally docked portions of the α-chain α220-249 and α412-472 in the model from Klykov *et al*. (PDBdev ID: 0000030) were removed to be extended with the α220-265 and α423-487 regions (renamed α200 and α400 hereafter, Supplementary Data 1) extracted from the AlphaFold2 model (downloaded from https://alphafold.ebi.ac.uk/entry/P02671) (Supplementary Fig. 1). The remaining dimer of trimers containing the structurally resolved part of the fibrinogen α, β and γ is hereafter defined as αβγ-main. Automatic detection of homodimers in XMAS, by means of storing separately overlapping peptide pairs, detected 10 reproducible overlapping peptide pairs from DSSO and DSS datasets as well as 4 in the PhoX dataset for the α400 domain only (Supplementary Data 2). In addition to these highly reproducible crosslinks, 1 self-link (α463-α463) was found in all datasets, confirming that the α400 region is a pivotal region in aggregating fibrinogen protofibrils irrespective of the sample source (plasma vs. commercial purified fibrinogen) and the sample preparation procedures (i.e. this work vs. Klykov *et al*. (33)). The detected distance constraints specifically defining the homodimer interaction interface were directly used as ambiguous interaction restraints (AIR) for the molecular docking using the HADDOCK 2.4 server (24). The AIR were submitted as a tbl file defining a median distance of 10 Å (lower bound up to 5 Å and upper bound up to 25 Å). The tbl file was generated from the XMAS environment as described. Representative structures of the top three clusters in the HADDOCK output were evaluated by mapping the submitted XL-MS restraints to select the model best supported by the experimental evidences (Fig. 2). A summary of the HADDOCK results, the parameter file, the distance constraints used as pseudobonds files and the top three clusters used in Fig. 2 are supplied in the “Supplementary Data 2” folder. The predicted model of the α400-dimer was then first used to dock the additional α200 to it (Supplementary Data 4), and subsequently the resulting α200-400 module to the main αβγ-core fibril (Supplementary Data 5). The preparation and interpretation of rotational sampling step using the DisVis server was performed as described in the section, “Integrative modeling in XMAS”. The “true positive” constraints were selected if their z-score was lower than 0.5 for the α200 to α400-dimer docking (Supplementary Data 4). A more stringent cut-off of 0 (*i.e*. only negative z-score values) was applied for the α200-400 module to the main αβγ-core fibril docking, given the much larger extent of the constraints considered, thus ultimately improving the precision of the subsequent docking step (Supplementary Data 5, Supplementary Figure 2). It should be carefully taken into account that, especially for the system here considered, XL-MS datasets may include (and often do) multiple conformations and/or interaction interfaces. What is here called a “false” positive, although reproducibly identified in independent experimental replicates, is therefore nothing more than a restraint incompatible with the current filtering method. These excluded restraints can be investigated separately following the same workflow here described, and possibly lead to define equally well supported interaction interfaces. For the subsequent molecular docking in HADDOCK, the multiple chains of the “fixed” chains used for DisVis rotational sampling (i.e. the α400-dimer and αβγ-core fibril) were merged into a single chain renamed as “A” and amino acid positions renumbered from “1” using PDBtools (34). Consequently the “scanning” chain, i.e. the structure we want to dock to the main “fixed” chain, was also renamed as “A” and amino acid positions renumbered from “1” using the same procedure. Those pseudobonds that were selected based on the z-score values (Supplementary Fig. 2) were then used to generate the AIR tbl restraint file with defined distances (i.e. 5-25 Å for PhoX or 6-33 Å for DSS). The models resulting from the HADDOCK runs were then re-evaluated in XMAS by mapping the XL-MS data on the best scoring HADDOCK clusters and comparing them based on XL-MS distances (top three in the examples for Fig. 2 and Fig. 3).

## Results

### Human fibrin clots as a model system

To demonstrate the capabilities implemented in XMAS, we used fibrin clots as our model system. Fibrin clots are formed by activation of fibrinogen hexamers, major constituents of plasma that assist in wound healing. Activation releases fibrinopeptides from fibrinogen, forming fibrin, after which clot formation occurs through oligomerization, followed by lateral aggregation, packing into fibrin fibers, and consequent branching (35). Fibrin(ogen) consists of three chains (α, β, and γ), for which the α-chain sports largely unstructured and flexible regions that are challenging to investigate by conventional structural biology approaches. Recently, XL-MS using DSSO crosslinking in combination with structural modeling has proven successful in unveiling structural details of fibrin clots formed in human plasma (33). To extend this dataset, we performed additional experiments on commercially available fibrinogen purified from human plasma for which a significantly lower amount (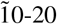 fold less) was clotted in triplicate independent reactions, and crosslinked with the two different reagents PhoX and DSS (see Materials and Methods for details). This dataset consists of 339 PhoX crosslinks (156 inter- and 183 intralinks) and 273 DSS crosslinks (122 inter- and 151 intralinks). The main goal is to use XMAS to extend the current protein model by taking advantage of the AlphaFold2 prediction for the α-chain (downloaded from https://alphafold.ebi.ac.uk/entry/P02671). From this model, we extracted the α220-265 and α423-487 (Supplementary Fig. 1 and Supplementary Data 1) domains (hereafter α200 and α400, respectively) in ChimeraX, and used the XMAS interface to DisVis/HADDOCK to integrate these protein sections into the current fibrin model. Although this increases the sequence coverage of the α-chain model by only a modest 4% with respect to the previous model, the extension provides structural details important for understanding lateral aggregation and cellular binding. It is worth noting that only those structural domains for which an average confidence was achieved (*i.e*. pLDDT > 50) and for which crosslinks have been identified were taken into account. In the following paragraphs, we use this biologically relevant system as a proof-of-concept to describe how this was achieved with XMAS, including how to detect homomeric interactions, and visualize, compare and analyze XL-MS data.

### Mapping Crosslinks to proteins structures, more than a simple visualization step

One of the major bottlenecks in the use of XL-MS derived distance constraints was the need to prepare results for loading into programs like ChimeraX through file re-formatting. To prevent the required labor intensive and error-prone file manipulations, XMAS natively reads the result files from a number of commonly used XL-MS search engines: XlinkX (36), pLink (37), and Xi (38). With its open architecture, more output types can easily be added to XMAS as well. Besides these raw outputs, XMAS includes support to the open standard file format mzIdentML, which is evolving to become a community standard for XL-MS data (39). To further improve user friendliness, XMAS performs automatic sequence alignment of the peptide sequences of the identified crosslinks (extracted from “Molecular models” in Fig. 1A) against the protein sequence(s) of the selected PDB structure(s) (“Evidence files” in Fig. 1A). This effectively prevents localization mistakes of the distance constraints in the structure. If a match is found for both crosslinked amino acids, a graphical object connecting their alpha-carbon atoms named pseudobond (or *pb*) is generated. After this step, the “Crosslink models” panel will then contain all the *pb* files generated with an evidence file for the given structures (Molecular models). Information on this mapping stage is provided in the “Log” window, including the location of a report file in tsv format where the *pb* strings and other metadata relevant to each crosslink of the initial evidence file. During the mapping stage, all possible combinations of residue-to-residue contacts are made if a chain is present multiple times in the selected models or the protein contains multiple stretches with the same sequence. This obviously generates too many pseudobonds mapped within the same structure, an issue which is resolved in a later step, but is nonetheless needed to avoid biases in the interpretation.

**Figure 1.**
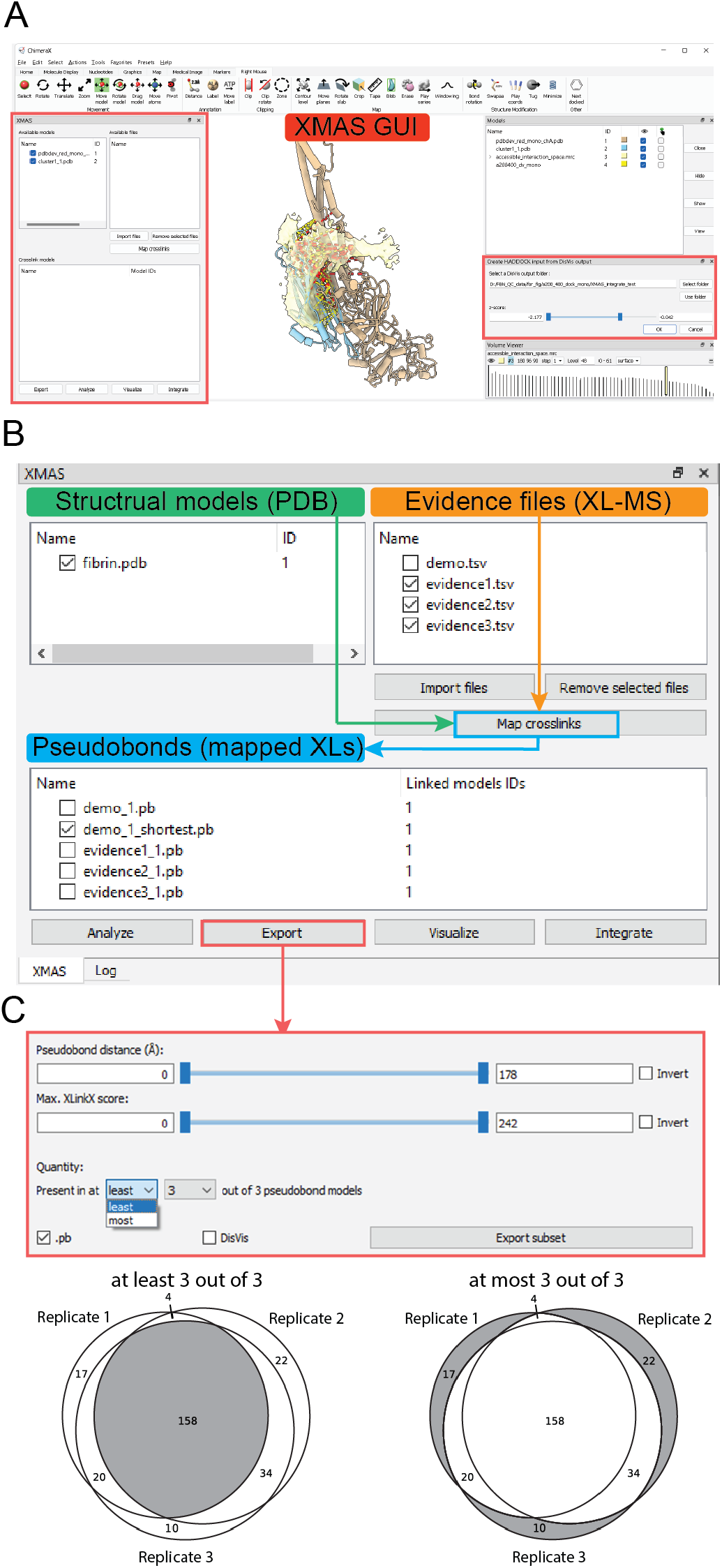
XMAS graphical user interface integrated in ChimeraX 1.3. Main window of ChimeraX 1.3 with localization of XMAS GUI **(A)**. The main XMAS interface listing available models (PDB) and evidence files (Crosslinks) that can be matched together with the button “Map Crosslinks” to generate a unique pseudobond file from one or more evidence file(s) with one or more available models (Model ID column, listed and identified by their respective numerical ID) **(B)**. As an example of the “Export” GUI functionalities, the selection of reproducible crosslinks comparing pseudobond models in at “least” (common to all datasets) or at “most” (uniquely represented in one or more datasets) **(C)**.

### Visualization, analysis and comparison

High-quality rendering of images and animations is a critical ability to communicate novel scientific discoveries effectively. ChimeraX maintains the ability to do so for proteins and protein complexes in stunning detail. To extend on its native capabilities, we added a “Visualize” menu to XMAS that unlocks capabilities to store and display useful data for each crosslink (i.e. pseudobond). Intuitively, the length of the pseudobonds represents the most valuable piece of information. In the XMAS GUI, it is possible to change the color and style of the pseudobonds according to an arbitrary crosslinker-related cutoff or the distance combined with a color gradient visualization (see Fig. 4 for examples). Next to this, XMAS can load multiple datasets, locate all unique residue pairs, and map all unique pairs onto the protein structures. Importantly, XMAS offers the option to visualize the overlap between replicates in a Venn diagram, providing critical insights into the success of the experiment (Fig. 1C). Additionally, we have over the years found it effective to also filter the datasets on reproducible links. XMAS offers the option to do so with a setting to select crosslinks detected in *n* out of the total number of experiments (Fig. 1C).

### Detecting homomeric interactions

During the mapping stage, XMAS also analyzes the identified peptide pairs to detect possibly occurring homomeric interactions. To achieve this, XMAS highlights those identifications where either the same residue (or position within the protein sequence) in a peptide pair is crosslinked to itself on the other peptide or where the two peptides share part of their sequence (Fig. 2A). As these options are physically impossible on the same protein (*i.e*. intra-links), such identifications can only arise from interacting homomers and this information can be used to drive the homomer modeling and validate the predictions. We name these peptide pairs “self-links” and, advantageously, these types of links can also be detected in existing datasets as they do not require additional, elaborate experimental setups. Comparing the crosslinked fibrin clots from human plasma (DSSO, Klykov *et al*. 2020 in Fig. 2B) to the newly generated data on *in-vitro* clotted fibrinogen (DSS and PhoX in Fig. 2B) it is clear that these homomeric interactions of the α400 domain are reproducibly detected, suggesting a central role in interfibril interactions. Although we do not further investigate this role here, we do supply the modeling steps in the following paragraphs that lead to a credible model.

**Figure 2.**
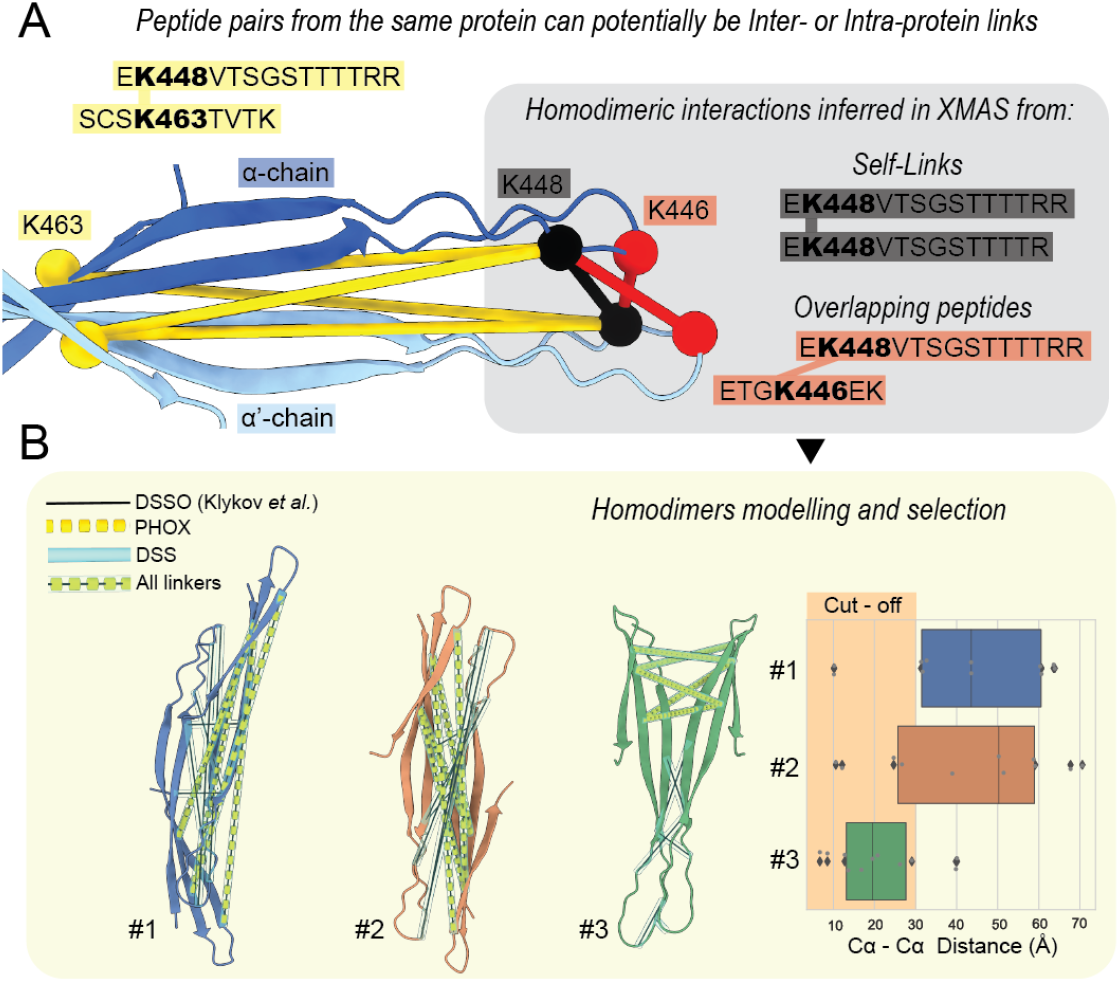
Unbiased mapping and detection of homomeric interactions. **(A)** Self-links and overlapping peptide pairs are automatically detected from the datasets and indicate positions where homomeric interaction interfaces are located. **FB)** After extracting these self-links through the XMAS user-interface, protein-protein docking software like HADDOCK can be used to assemble the homodimer, resulting in multiple models that can be validated/filtered with the crosslink restraints distances.

### Integrative modeling assisted by XMAS

With credible structures in hand, for which we here use the existing αβγ-core fibril model and the AlphaFold2 predicted versions for the α400 dimer and α200, the current fibrin model can be extended to cover more of the amino acid sequence. To achieve this, all the structures can be loaded into a single ChimeraX session and our XL-MS distance constraints mapped, after which we aim to uncover the “inter-model” crosslinks. As the next step, a selection needs to be made of the subset of XL-MS derived distance constraints that are consistent with a certain position of the α400-200 module in relation to the αβγ-core fibril. For this, we rely on the DisVis package (31), where the αβγ-core fibril model is viewed as “fixed” and the α400-200 module as ”scanning” chain. The required input files are interactively generated in XMAS from the “Export” menu (Fig.1B) where the “scanning” and “fixed” chains, the crosslinks datasets to include, and the minimum/maximum distance specific for the crosslinker in use (*e.g*. 5-25 Å for PhoX, 6-33 for DSS and DSSO, *etc.*) can be set. The resulting TXT file containing the distance constraints together with the two PDB structures can be uploaded to the freely accessible at https://wenmr.science.uu.nl/disvis/. When the run has completed, the DisVis output folder can directly be opened through the “Integrate” menu. XMAS will load both structures, open the MRC density showing the maximum number of consistent constraints at every position in space, and all the input constraints mapped as a new set of pseudobonds (Fig. 3). As ChimeraX allows the user to rotate and move the “scanning” model, XMAS also color-codes the pseudobonds that either violate (Red) or agree with (Yellow) the distance defined in the DisVis input file (Fig. 3). For the selection of which consistent constraints to keep for a subsequent molecular docking step in HADDOCK, an interactive slider selector for the z-scores is supplied where higher scores denote a more likely “false positive” distance constraint (in DisVis parlor). It should be noted, however, that in some cases more than one interaction interface might be defined in the occupancy analysis (e.g. if a protein has multiple binding sites) and therefore constraints excluded merely based on a z-score cut-off may not be “false positives”, but simply point to another position of binding that is supported by fewer crosslinks. When selecting *Integrate* → *Create HADDOCK input*, the desired distance constraints defining the interaction are then exported to a TBL file (formatted in the CNS syntax that is accepted by HADDOCK) as wel as the fixed chain as ChainA.pdb and the scanning chain as ChainB.pdb to be directly uploaded at https://wenmr.science.uu.nl/haddock2.4 for the actual docking of the two molecules. When the standard settings apply, these files are the only ones that need to be set in the extensive set of options available for HADDOCK (the TBL file as unambiguous restraints). After a successful docking run, the clustered solutions can be evaluated in light of the XL-MS experimental evidences and the better predicted models selected based on the detected distance constraints (Fig. 3B).

**Figure 3.**
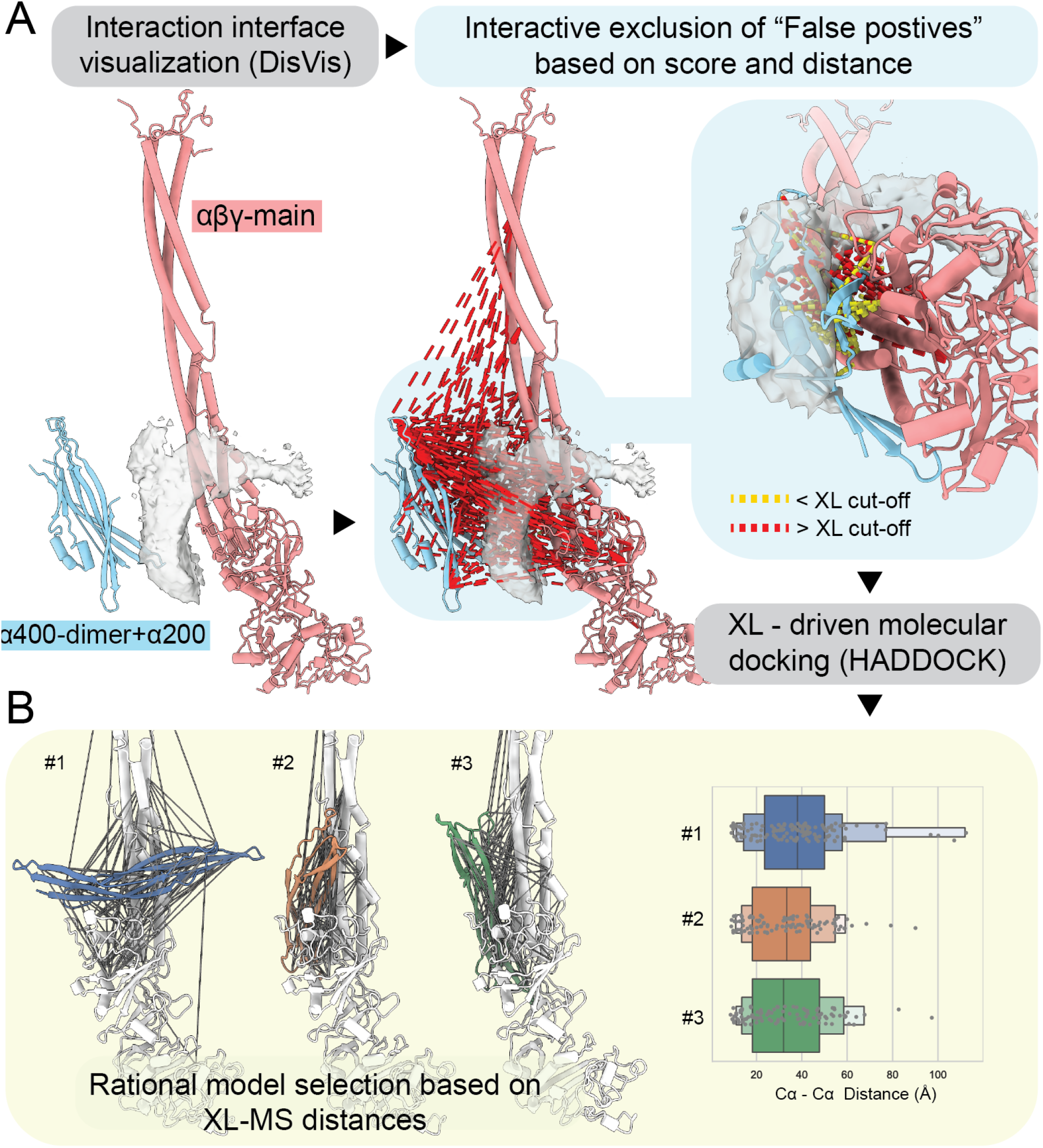
XMAS assisted integrative modeling. **(A)** The location of novel protein stretches or interacting proteins can be located based on the distance constraints and DisVis – a process assisted by XMAS in obtaining the desired distance constraints from the complete set and producing the required input files for DisVis. **(B)** After further filtering the distance constraints in XMAS based on the DisVis results, the input files for HADDOCK are generated from XMAS and the docking run can be started and outputs evaluated in the context of experimentally derived distance constraints

### XMAS visualization capabilities allow a deeper analysis and comparison of large datasets

Once the integrative model is generated, XMAS aims at integrating advanced options for high-quality rendering of images and animations. This is accomplished thanks to the integration into ChimeraX, allowing the graceful and smooth visual handling of new and large data types. We added the “Visualize” menu, where capabilities to store and display useful data for each crosslink (i.e. pseudobond) are located. Intuitively, the length of the pseudobonds represents the most valuable piece of information. In the XMAS GUI it is possible to change the color and style of the pseudobonds according to an arbitrary crosslinker-related cutoff that can eventually be combined with a color gradient visualization for the distance (Fig. 4). Besides applications of XL-MS on purified or recombinantly expressed protein complexes, one of the strengths of this technique is its applicability at systems-wide scales (18, 40, 41). The integration of XMAS in the ChimeraX environment additionally eases the handling of large structures and/or the many structures uncovered by systems-wide XL-MS datasets. This is demonstrated here for a publicly available dataset (PXD008418) generated from a crosslinking experiment on PC9 cell lysate yielding >2000 crosslinks (25). In this example, we uploaded the PDB codes of the top 18 proteins with > 1 crosslink directly from the [PDB databank](https://www.rcsb.org/) and mapped the respective crosslinks with the “find shortest” option. The result is a rational-based rapid evaluation of the proteome-wide experiment quality (Supplementary Fig. 2).

**Figure 4.**
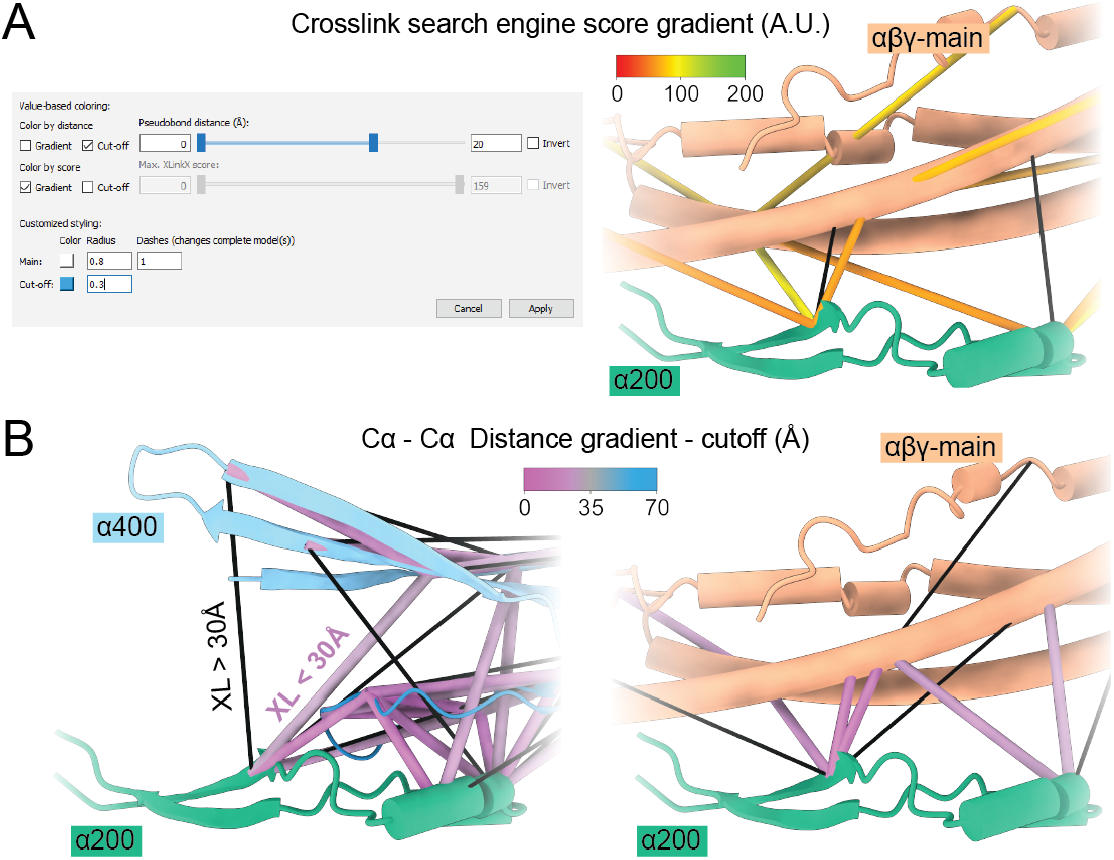
Advanced visualization options supported by XMAS. **(A)** Various settings pertaining to the distance constraints can be made (left panel), with the identification score as example (right panel). **(B)** The distance spanned by the distance constraints can easily be colored.

### Future developments, usability and community support

The open architecture of XMAS, its distribution as open-access software hosted at GitHub and the possibility to install it from the ChimeraX toolshed will allow non-expert users to use the tool and expert users to build up novel and tailored functionalities in future releases. From our side, future features will include: 1) dedicated pipelines for the visualization and analysis of quantitative data, 2) direct visualization of annotated MS spectral data, and 3) integration with Molecular Dynamics simulation trajectories to analyze protein conformational changes. With the ease XMAS brings to structural modeling with XL-MS data, we anticipate more projects to leverage this type of data. The combination of Cryo-electron Tomography with XL-MS and protein computational modeling will also likely represent a step toward the visualization of the molecular sociology of the cell, for which XMAS already provides ample tools (42).

## Conclusions

The field of XL-MS is rapidly expanding and gaining popularity among structural and cell biologists. The visualization and interpretation of these data in a structural context are however relying on piecemeal strategies, mainly implemented in specialized massspectrometry or molecular modeling-oriented laboratories. To alleviate this and to popularize the use (and re-use) of XL-MS data in an open fashion, we designed XMAS to fulfill the needs of both proficient and inexperienced users. The key features provided are 1) even further simplifying the connection to HADDOCK, 2) easy loading of crosslinks regardless of the used search engine and without the need for data (re)formatting, 3) a robust mapping and visualization procedure relying on sequence alignment directly into PDB structures and detecting homodimeric interactions based on peptide sequence overlap, 4) analytical and plotting capabilities resulting in publication-ready figures and plots, 5) integration with EVouplings(43), and 6) a seamless interfacing with integrative modeling tools other than HADDOCK. XMAS is integrated into the fantastic ChimeraX environment, with its many user-friendly and stunning visualization options. With these features on board, we think XMAS provides a future proof option for the integration of XL-MS data in structural modeling projects.

## Supporting information

Manual

Practice dataset

## Acknowledgments

The Dutch Research Council (NWO) supported this research through funding of the large-scale proteomics facility Proteins@Work (project 184.032.201) embedded in the Netherlands Proteomics Centre. We further acknowledge funding through the European Union Horizon 2020 program INFRAIA project Epic-XS (Project 823839). Finally, this work is part of the research program NWO TA with project number 741.018.201, which is partly financed by NWO.

## Contributions

IL: implementation, software design, writing; PA: conceptualization, design, supervision, writing; supervision; AJ: implementation; RAS: conceptualization, funding acquisition, supervision, writing.

## Declaration of interests

The authors declare no competing interests.

## Supplementary Material

In the supplementary material two files are uploaded that contain all material for the manual. The file *20220421_manual.pdf* contains a complete description of the package with all steps described in detail. The file *20220421 _example_data.zip* contains example data that can be used in conjunction with the manual to practice all functionality.

Appended to this manuscript are the supplementary figures.

## Data availability

The mass spectrometry proteomics data have been deposited to the ProteomeXchange Consortium via the PRIDE partner repository with the dataset identifier PXD033409.

---

**Supplementary Figure 1:**
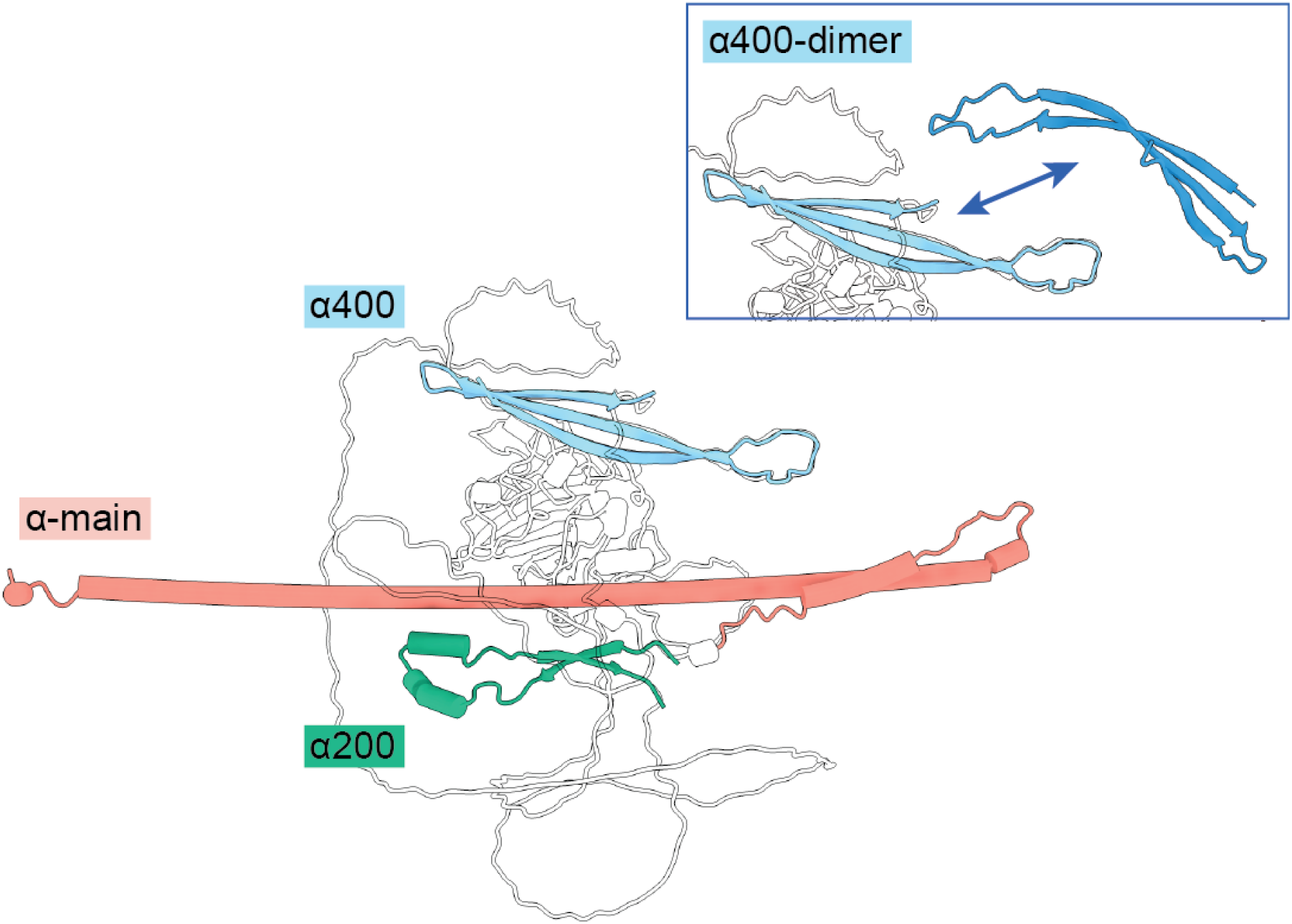
Individual structures used for Integrative modeling of the Fibrin chain. The α200 (α220-265) and α400 (α423-487) regions extracted from the AlpahFold2 model (downloaded from https://alphafold.ebi.ac.uk/entry/P02671) of the Human fibrinogen α chain (UniProt ID: P02671). These models were used in this work in combination with newly acquired XL-MS data and XMAS to extend the previously available structural model.

**Supplementary Figure 2:**
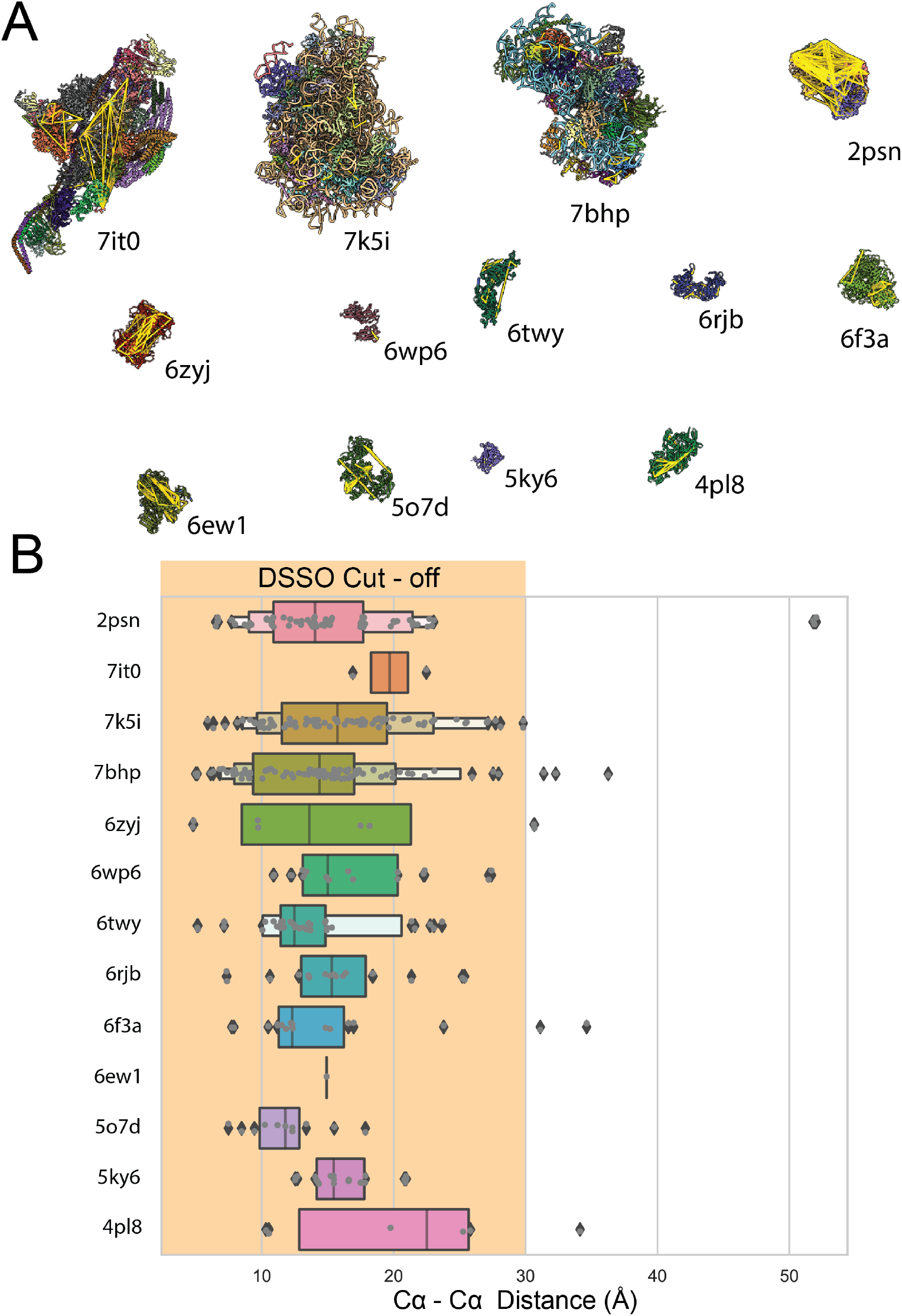
Evaluation of proteome-wide XL-MS experiment quality. XMAS integrated into ChimeraX allows the smooth handling of multiple large structures and datasets. In this example, 13 PDB structures (PDB IDs indicated) were selected from the list of the one with most identified crosslinks from a proteome-wide XL-MS experiment on human cell lysate (26). Crosslinks can be mapped in all structures within seconds **(A)** and plotting the distance of the “shortest” for each peptide pair, to avoid biases due to homomeric complexes, allows for a rapid and throughout evaluation of the data quality **(B)**.

**Supplementary Figure 3:**
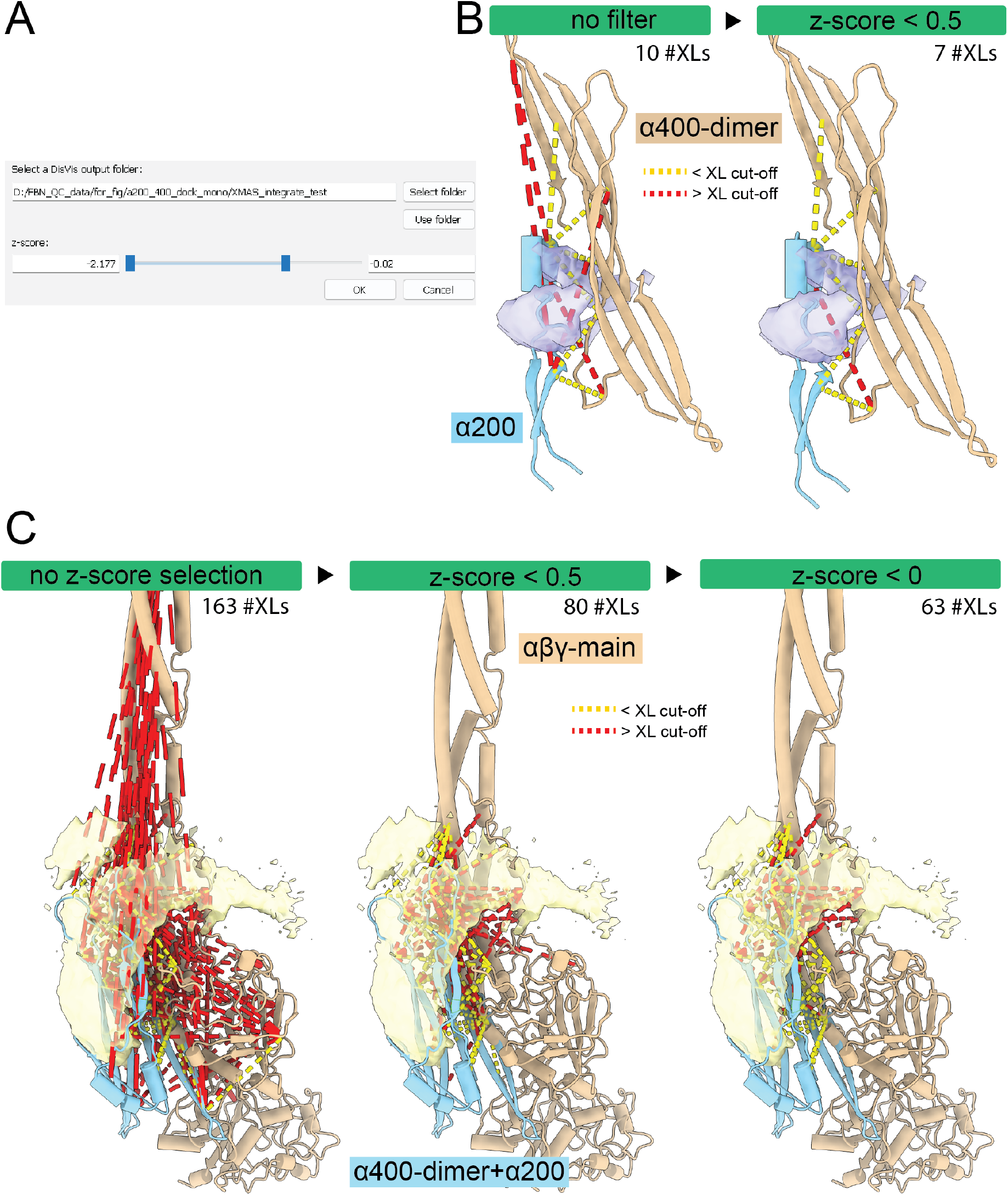
Examples of the integrative modeling pipeline through XMAS implemented in this work. The “integrate” XMAS module was used to filter out putative false positive restraints determined by DisVis using the z-score selector **(A)**. The valid restraints according to the selected cut-off are automatically appearing/disappearing while the slider moves, allowing the user to evaluate the data in real-time. In addition, color coding based on the distance cutoff set in the DisVis restraint file is automatically read and applied (in this case pseudobonds <5 Å or >25 Å are colored in red). In these two examples the number of selected crosslinks further used for docking the α200 to α400 dimer **(B)** and the resulting α400dimer + α200 module to the αβγ-main chain **(C)** are indicated for 2 and 3 steps of z-score cut-off, respectively.

## Notes

https://cxtoolshed.rbvi.ucsf.edu/apps/chimeraxxmas

### Competing Interest Statement

The authors have declared no competing interest.

### Summary of Updates

We added full support for HADDOCK protein-protein docking to the software. The paper was updated accordingly.

