## Supplementary material for "Xlink Mapping and AnalySis (XMAS) - Smooth Integrative Modeling in ChimeraX": Manual

### XMAS User Manual

Scheltema Lab, 21 April 2022

https://github.com/ScheltemaLab

### 1. About this manual

This manual explains how to use XL Mapping and AnalySis (XMAS), a tool for downstream analysis and visualization of crosslinking mass spectrometry (XL-MS) data. XMAS is a bundle for the molecular visualization software UCSF ChimeraX (hereafter called ChimeraX)[1]. The information about ChimeraX that is required to understand this manual is provided. For further information about ChimeraX, please visit the ChimeraX web page.

### 2. Dictionary

- **Accessible interaction space** (DisVis) – area representing the collection of the centers-of-mass of all scanning chain positions for a given number of consistent restrains.
- **Atom name** – name used in ChimeraX to refer to a type of atom.
- **Atom specifier string** – set of characters used to refer to an atom in a ChimeraX session. Atom specifier strings are formatted as follows: #*w*/*x*:*y*@*z*, where *w* represents the model ID, *x* represents the chain ID, *y* represents the residue number, and *z* represents the atom name.
- **Available set** (pseudobonds) – regarding XMAS's Analyze, Export, and Visualize windows, the available set refers to the pseudobonds that were selected at the moment that the button to open the window was clicked.
- **Chemical linker** – in crosslinking mass spectrometry, a chemical linker is a compound that harbors the potential to covalently bind to two protein residues, thereby linking these residues to each other.
- **Confidence score** – numerical value reflecting the likelihood that an identified crosslink is a true positive. In the evidence files, the confidence scores for XlinkX, Xi, pLink, and mzIdentML can be found under the 'Max. XlinkX Score' column, 'Score' column, 'Score' column, and 'xi:score' tag, respectively.
- **Consistent restraint** (DisVis) – a crosslink with a distance that is consistent with the distance restraint.
- **Crosslink** – two peptides, covalently linked by a chemical linker.
- **Crosslink models** – graphical models representing a collection of crosslinks.
- **Distance** (crosslink/pseudobond) – the length of the crosslink or pseudobond.
- **Distance restraint** – known distance or range of distances between two atoms.
- **DisVis** – webserver for analyzing the information content of distance restraints between two molecular models.
- **Evidence file** – file containing identified crosslinks. From this file, XMAS uses the peptide amino acid sequences, positions of the crosslinked residues, and confidence scores.
- **Fixed chain** (DisVis) – term assigned to one of the two molecular models in a DisVis job. The other model is called the scanning chain.
- **ID**
    - **Chain ID** – unique alphabetical code that refers to a chain in a ChimeraX model.
    - **Model ID** – unique numeric code that refers to a ChimeraX model open in the session.
- **Interlink** – crosslink connecting residues belonging to different proteins.
- **Interlink** (XMAS)
    - **Chain interlink** – PB connecting atoms that belong to different chains of the same ChimeraX model.
    - **Model interlink** – PB connecting atoms that belong to different ChimeraX models.
- **Intralink** – crosslink connecting residues belonging to the same protein.
- **Intralink** (XMAS) – PB connecting two atoms in the same ChimeraX chain.

- **Linked model** – model that contains one or more crosslinked residues; each crosslink model has one or more linked models.
- **Mapping** – aligning the peptide amino acid sequences from peptide pairs in evidence files to the amino acid sequences of molecular models, and thereby determining possible positions of crosslinks.
- **Mapping information file** – tab-delimited file (extension '.tsv') generated for each evidence file that was mapped with XMAS, containing mapping results for each peptide pair.
- **Molecular models** – graphical models representing molecules.
- **Overlap associated** – when the alignments of both peptides of a peptide pair on a molecular model do not overlap, but combinations of overlapping alignments have been found for the particular peptide pair.
- **Overlapping (non-self)** – when the sequence alignments of both peptides of a peptide pair on a molecular model overlap in such a way that the crosslinked residues of the peptides are different, as opposed to identical (as in the overlapping (self) category).
- **Overlapping (self)** – when the sequence alignments of both peptides of a peptide pair on a molecular model overlap in such a way that the crosslinked residues of the peptides are identical, as opposed to different (as in the overlapping (non-self) category).
- **Peptide pair** – the two peptides in a crosslink.
- **pLink** – search engine to identify crosslinks from experimental crosslinking mass spectrometry data.
- **Proteome Discoverer** – search engine to identify crosslinks from experimental crosslinking mass spectrometry data.
- **Pseudobond** – graphical component in ChimeraX, visualizing the shortest possible trajectory between two specific atoms as a line (often dashed).
- **Pseudobond file** – ChimeraX readable file (extension '.pb') to save pseudobonds and reopen them as a pseudobonds model in a ChimeraX session.
- **Pseudobond specifier string** – the atom specifier strings of the two atoms that are connected by a pseudobond, separated by a space.
- **Residue number** – the index of a residue in the amino acid sequence of a ChimeraX chain.
- **Restraints file** (DisVis) – file (extension '.txt') containing the PBs and the distance restraints applicable to them that can be used as input in a DisVis job.
- **Scanning chain** (DisVis) – term assigned to one of the two molecular models in a DisVis job. The other model is called the fixed chain.
- **Selflink** – theoretical situation in which a chemical linker attaches to the same residue twice.
- **Xi** – search engine to identify crosslinks from experimental crosslinking mass spectrometry data.
- **XlinkX** – node that can be incorporated into Proteome Discoverer, to identify crosslinks from experimental crosslinking mass spectrometry data.
- **Z-score** (DisVis) – score reflecting the likelihood that a restraint is a false-positive.

#### 3. Abbreviations

- **Å** – Ångström
- **PB** – pseudobond
- **PD** – Proteome Discoverer
- **PDB** – Protein Data Bank
- **XL-MS** – crosslinking mass spectrometry
- **XMAS** – XL-MS Mapping and AnalySis

#### 4. Introduction to XL-MS

In XL-MS, protein residues are covalently linked, or 'crosslinked' together by adding chemical linkers (**Figure 1**). Since chemical linkers with known lengths are used, this provides distance restraints between the crosslinked residues, yielding structural information. To obtain the distance restraints, the crosslinked proteins are digested, yielding short fragments called peptides. Since the covalent chemical linkage remains intact during the digestion, part of the peptides are crosslinked to other peptides[2,3]. A chemical linker connecting two peptides is called a crosslink, and the connected peptides are called a peptide pair[2]. After the digestion, samples are measured using mass spectrometry. Subsequently, the resulting spectra are analyzed with a search engine to identify the crosslinks. For each of the peptides in a peptide pair, the amino acid sequence and the crosslinked residue (i.e. the residue that the chemical linker is attached to) are determined. Thereby, the location of each crosslinked residue in its protein is found. Crosslinks within a protein, called intralinks, provide information on tertiary structure, whereas crosslinks between proteins, called interlinks, provide information on the quaternary structure of protein complexes[2,3]. The search engine additionally calculates a confidence score for each crosslink[2-4].

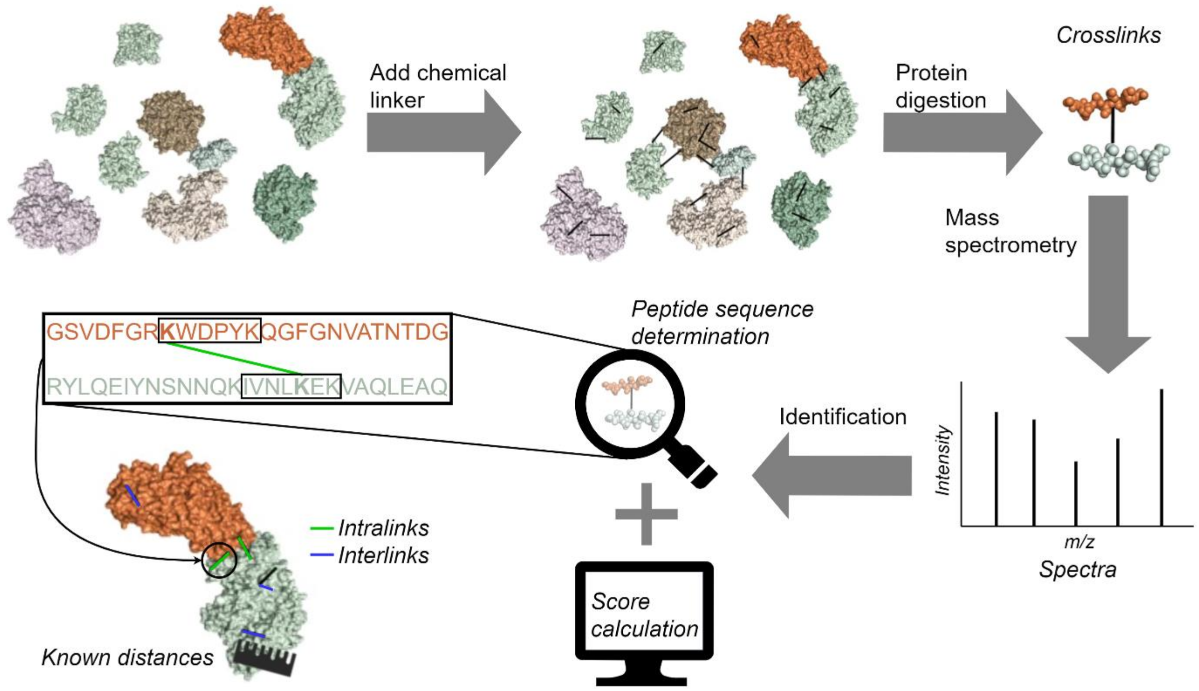

**Figure 1. Basic XL-MS workflow.**
Adapted from Klykov et al.[2].

#### 5. Setting up

To use XMAS, users must install ChimeraX and XMAS, and the XMAS main window must be opened. Changing the main window's position is optional.

##### 5.1. Installing ChimeraX

Before XMAS can be installed, ChimeraX (version 1.3 or higher) should be installed on the machine used. Instructions for installation of ChimeraX can be found here.

#### 5.2. Installing XMAS

XMAS is currently distributed as a folder, that can be downloaded from GitHub. One of the items in this folder is the file `INSTALL.txt`. Follow the instructions in `INSTALL.txt` to install XMAS on ChimeraX.

#### 5.3. Opening the main window

The XMAS main window can be opened in any ChimeraX session, using the ChimeraX Graphical User Interface (GUI) or command line. The general layout of the main window is shown in **Figure 2**.

##### 5.3.1. Opening using the GUI

In the menu bar, navigate to 'Tools' → 'Structure Analysis' → 'XMAS'.

##### 5.3.2. Opening using the command line

Command line: execute the command 'ui tool show XMAS'. Commands are executed by typing them in the ChimeraX command line interface and pressing 'Enter'.

#### 5.4. Positioning the main window

The XMAS main window can be positioned anywhere on screen using dragging and dropping. It can also be docked on all sides of the ChimeraX main window and on other tool windows.

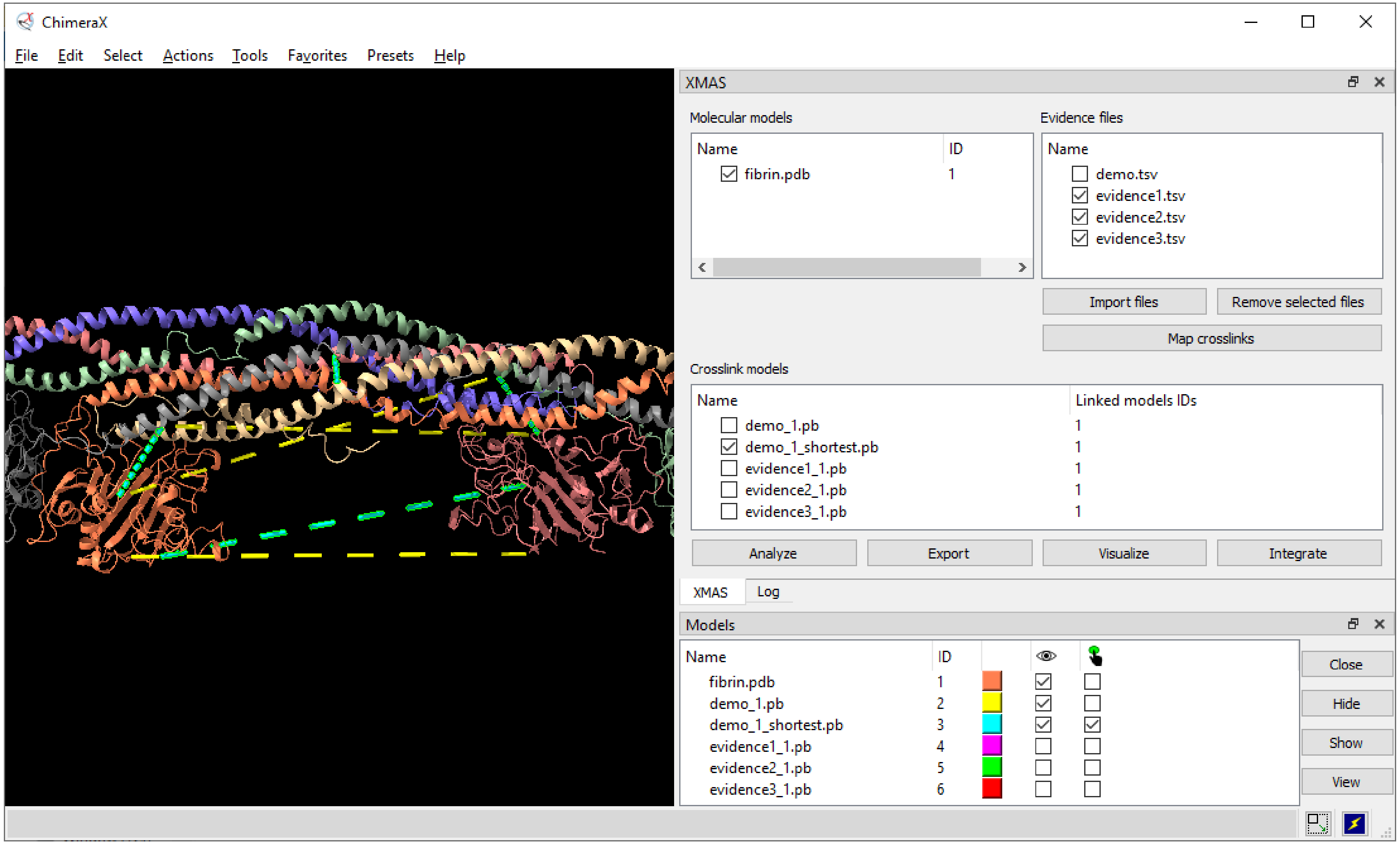

**Figure 2. Layout of the XMAS main window in a ChimeraX session.**

The ChimeraX Toolbar and Command Line Interface, tools that are shown by default, are hidden for clarity. XMAS is docked on the ChimeraX Log. Six models are open in the session: fibrin.pdb, demo_1.pb, demo_1_shortest.pb, evidence1_1.pb, evidence2_1.pb, and evidence3_1.pb. The first, fibrin.pdb, is a dimeric model of the protein fibrin. Since a fibrin monomer consists of three chains, the model consists of six chains, each colored distinctly. The remaining models are pseudobond models linked to fibrin.pdb. Only fibrin.pdb, demo_1.pb, and demo_1_shortest.pb are displayed; the other models are hidden. The selected model is demo_1_shortest.pb. Four files have been imported in XMAS: demo.tsv, evidence1.tsv, evidence2.tsv, and evidence3.tsv.

### 6. Managing models

XMAS interacts with ChimeraX to enable managing molecular and crosslink models in the XMAS model panels. This interaction involves opening and closing both models types, and selection of crosslink models.

#### 6.1. The XMAS model panels

The XMAS main window contains two model panels: the ‘Molecular models’ panel on the top left and the ‘Crosslink models’ panel below (**Figure 2**). These panels contain information on models relevant for XMAS; the ‘Molecular models’ panel contains the names and IDs of molecular models open in the ChimeraX session, and the ‘Crosslink models’ panel contains the names and ‘Linked models IDs’ of the crosslink models open in the session.

#### 6.2. Opening and closing models

When opening a molecular or crosslink model in the ChimeraX session, the model information automatically appears in the correct XMAS model panel. NB for a model to appear in ‘Crosslink models’, its name needs to end with ‘.pb’. Closing a model in the ChimeraX session removes it from its XMAS model panel.

#### 6.3. Selecting crosslink models

When the checkbox of a model in ‘Crosslink models’ is checked or unchecked, it is automatically selected in the ChimeraX session, and vice versa. The advantage of selecting in ‘Crosslink models’, as opposed to the ChimeraX ‘Models’ window, is the possibility to select multiple models simultaneously.

### 7. Managing evidence files

Clicking ‘Import files’ underneath the ‘Evidence files’ panel opens a dialog where one or more evidence files can be selected, which then appear in the panel. Files are removed from the panel by selecting their name (not the checkbox) and, subsequently, clicking ‘Remove selected files’.

XMAS currently supports evidence files generated with the search engines XlinkX for Proteome Discoverer (PD), Xi, and pLink, in csv, tsv, txt, and xlsx format. Additionally, mzIdentML files (mzid format) are supported.

### 8. Mapping crosslinks

XMAS enables mapping the crosslinks in any number of evidence files on any number of molecular models. For each evidence file, a mapping log and mapping information file are created. Moreover, if crosslinks could be mapped onto the structure(s), a crosslink model and PB file are generated.

#### 8.1. Creating crosslink models

To select the molecular models and evidence files to use for mapping, their checkboxes are checked in the ‘Molecular models’ and Evidence files’ panel, respectively. Clicking ‘Map crosslinks’ then initiates sequence alignment of all peptide pairs in all checked evidence files to all checked molecular models. If at least one perfect alignment has been found for each peptide in a peptide pair, pseudobonds (PBs) are generated; one PB for each combination of these perfect alignments. For more information, see 8.4. Interpreting the mapping information file. All PBs are grouped in PB models; one model for each mapped evidence file. The name of a PB model is equal to `*name_IDs*.pb`, where *name* represents the name of the evidence file, and *IDs* represent the ID or IDs of the model or models it was mapped to. Consequently, the PB model created by mapping `demo.tsv` on `fibrin.pdb` is called demo_1.pb, because the ID of fibrin.pdb is 1 (**Figure 2**). Created files (PB files and mapping information files) are stored in the same location as the evidence file. The file names are identical to the names of the corresponding PB model, where the '.pb' extension is replaced with '.tsv' for the mapping information file.

#### 8.2. Interpreting the mapping log

For each mapping job, a log is displayed in the ChimeraX 'Log' window (**Figure 3a**). The mapping log contains the following pieces of information (items marked with an asterisk (*) are only included when applicable):

- **Evidence file** – 'Peptide pair mapping of *engine* evidence file: *file*', where *engine* represents the search engine used to generate the evidence file mapped in the current mapping job, and *file* represents the path to that evidence file.
- ***Lacking/decoy sequence** – '*x* Peptide pairs are disregarded due to lacking/decoy sequence', where *x* represents the number of peptide pairs with a lacking or decoy peptide sequence. These peptide pairs are ignored in the mapping procedure. This remark is excluded if all peptides contained sequence information.
- **Unique peptide pairs** – 'Unique peptide pairs: *x* out of *y*', where *x* represents the number of unique peptide pairs, and *y* represents the total number of peptide pairs in the evidence file. The difference between both numbers equals the number of duplicate peptide pairs. No PBs are formed for these peptide pairs.
- **Peptide pairs with PBs** – 'Unique peptide pairs with pseudobonds: *x*', where *x* represents the number of peptide pairs for which PBs have been found.
- ***PB file** – 'Pseudobonds are stored in *file*', where *file* represents the path to the PB file generated for the current mapping job. This remark is only included if PBs were created in this mapping job.
- ***Mapping information file** – 'Mapping information is stored in *file*', where *file* represent the path to the mapping information file generated for the current mapping job. This remark is only included if PBs were created in this mapping job.

#### 8.3. Interpreting the PB file

The PB file is especially useful to open PBs in another session than the session in which they were created. Each line in the file is a PB specifier string, representing one PB (**Figure 3b-d**). XMAS connects the α carbon atoms of the crosslinked residues. Therefore, the atom names for the two atoms in the string is 'CA'; the ChimeraX code for α carbon. NB besides PB specifier strings, PB files might contain additional information. Firstly, comment lines starting with a semicolon might be present. Secondly, additional information can be included after the PB specifier. This additional information is explained in the ChimeraX User Guide.

#### 8.4. Interpreting the mapping information file

The mapping information file enables tracing back the mapping results to the peptide pairs in the evidence file. Each peptide pair in the evidence file gets one or more rows in the information file (**Figure 3e**). Each row has four columns: Reference to peptide pair, 'Pseudobond', 'Overlap category', and 'Distance (A)'.

##### 8.4.1. Reference to peptide pair

Column that refers back to the peptide pair in the evidence file. The type of reference depends on the evidence file format, and can be inferred from the column header. In XlinkX evidence files, the value in the reference column is equal to the row number in the evidence file. For Xi, pLink, and mzIdentML evidence files, the value refers to the 'PeptidePairID' or 'PSMID' column (depending on which one is present in the evidence file), the 'Peptide_Order' column, and the 'Peptide id' tag, respectively.

##### 8.4.2. 'Pseudobond'

Can contain four types of data:

- **'Sequence lacking'**, if peptide sequence information was lacking for this peptide pair in the evidence file.
- **'No pseudobonds'**, if no PBs were found for this peptide pair, even though peptide sequence information was available.
- **'Duplicate of *x*'**, where *x* represents the identical peptide pair.
- A **PB specifier string**, specifying a PB that belongs to this peptide pair. All PBs belonging to one peptide pair get one row, meaning that peptide pairs with multiple PBs will get multiple rows. For example, peptide pair 7 in **Figure 3e** has four PBs, since there is one PB for each of the four combination of chains; B to C, B to F, C to E, and E to F.

##### 8.4.3. 'Overlap category'

Can contain four types of data:

- **No data**, when no PBs were found for a peptide pair.
- **'Overlapping (non-self)'**, for PBs between two residues that result from overlapping alignments (**Figure 3f**). In reality, residues cannot be shared between the two peptides of a pair. Hence PBs in this category are excluded from the PB model. However, unlike PBs of the 'overlapping (self)' category, 'Overlapping (non-self)' PBs can be opened in a ChimeraX session by storing their specifier strings in a PB file and opening this file in ChimeraX.
- **'Overlapping (self)'**, for PBs that result from overlapping alignments and that attach to the same residue twice, i.e. selflinks (**Figure 3f**). Like PBs of the 'overlapping (non-self) category, these PBs are excluded from the PB model, because selflinks do not occur in reality. Since ChimeraX does not support formation of a PB that links an atom to itself, it is impossible to open PBs of the 'overlapping (self)' category in ChimeraX.
- **'Overlap associated'**, when the PB belongs to a peptide pair for which overlapping alignments were found, but the PB itself does not result from overlapping alignments (**Figure 3f**). For example, peptide pair 5 in **Figure 3e** has two PBs from overlapping alignments, where both peptides are aligned on chain B, or both on chain E. However, this peptide pair also has two PBs from non-overlapping alignments, where one peptide is aligned on chain B, and the other on chain E, and vice versa. Given that peptide pair 5 has both overlapping and non-overlapping alignments, the PBs from non-overlapping alignments are categorized as 'Overlap associated'.
- **'Not overlap associated'**, when the PB belongs to a peptide pair for which no overlapping alignments were found (**Figure 3f**).

##### 8.4.4. 'Distance (A)'

The distance between the atoms linked by a PB at the moment that 'Map crosslinks' was clicked, in Ångström (Å). Distances are only shown for non-overlapping PBs, since only these are included in the PB model. The distance column can be updated.

**a**

Log

**Peptide pair mapping of XlinkX evidence file: C:/demo.tsv**
1 Peptide pair is disregarded due to lacking/decoy sequence
Unique peptide pairs: 4 out of 5
Unique peptide pairs with pseudobonds: 3
Pseudobonds are stored in C:/demo_1.pb
Mapping information is stored in C:/demo_1.tsv

**b**

1. #1/A:142@CA #1/D:142@CA
2. #1/B:295@CA #1/C:151@CA
3. #1/B:295@CA #1/F:151@CA
4. #1/B:367@CA #1/E:374@CA
5. #1/B:374@CA #1/E:367@CA
6. #1/C:151@CA #1/E:295@CA
7. #1/E:295@CA #1/F:151@CA

**c**

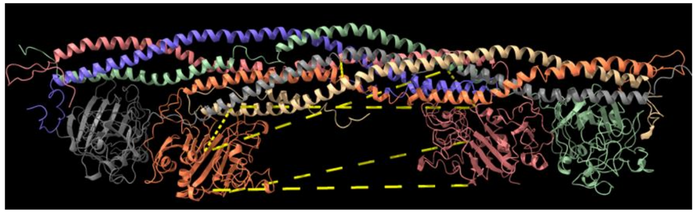

**d**

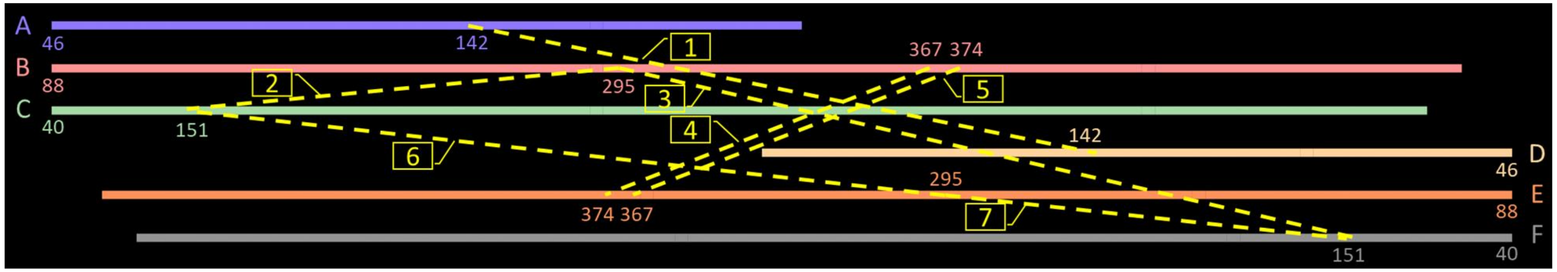

**e**

| Row in evidence file | Pseudobond | Overlap category | Distance (Å) |
|---|---|---|---|
| 2 | Sequence lacking | | |
| 3 | No pseudobonds | | |
| 4 | Duplicate of 3 | | |
| 5 | #1/B:367@CA #1/B:374@CA | Overlapping (non-self) | |
| 5 | #1/B:367@CA #1/E:374@CA | Overlap associated | 102.6567235 |
| 5 | #1/B:374@CA #1/E:367@CA | Overlap associated | 103.9982904 |
| 5 | #1/E:367@CA #1/E:374@CA | Overlapping (non-self) | |
| 6 | #1/A:142@CA #1/A:142@CA | Overlapping (self) | |
| 6 | #1/A:142@CA #1/D:142@CA | Overlap associated | 39.13762494 |
| 6 | #1/D:142@CA #1/D:142@CA | Overlapping (self) | |
| 7 | #1/B:295@CA #1/C:151@CA | Not overlap associated | 19.99902553 |
| 7 | #1/B:295@CA #1/F:151@CA | Not overlap associated | 98.65143901 |
| 7 | #1/C:151@CA #1/E:295@CA | Not overlap associated | 98.82706537 |
| 7 | #1/E:295@CA #1/F:151@CA | Not overlap associated | 20.47320859 |

**f**

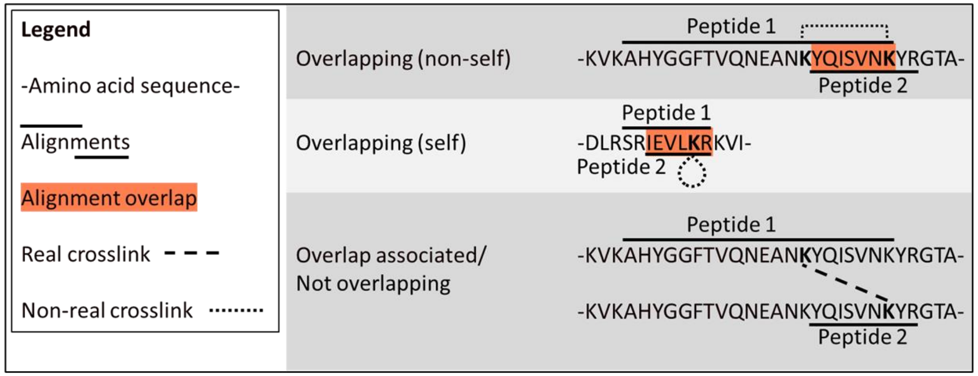

**Figure 3. Mapping job of an XlinkX evidence file on a molecular model.**

**a.** The mapping log. **b.** Lines in the generated PB file. Note that the line numbers are not present in actual PB files. **c.** The molecular model with the generated PB model in ChimeraX. **d.** Schematic representation of **b** and **c**. The six chains from the molecular model in **c** are depicted as continues lines with corresponding colors. Chains A-C are shown in a left-to-right orientation, and chains D-F in a right-to-left orientation. Chain IDs and the first residue numbers are depicted at the start of each chain. PBs are displayed as yellow dashed lines between chain residues, of which the residue numbers are visualized. The numerical labels on the PBs correspond to the line numbers in **b**. **e.** The mapping information file. **f.** Schematic examples of the three overlap categories in **e**.

### 9. Analyzing crosslinks

To analyze a set of PBs (the available set), these PBs are selected and the 'Analyze' button in the XMAS main window is clicked, which opens the Analyze window (**Figure 4a**). This window provides four functions: plotting overlap between multiple PB models, plotting distances, finding the shortest PBs per peptide pair, and updating distances in the mapping information file. Moreover, the Analyze window enables customizing the names used for plot labels and generated models. Created plots can be saved by clicking the save icon 🖫 in the plot window.

#### 9.1. Plotting overlap

Clicking 'Plot overlap' visualizes the overlap between PB models in a Venn diagram. A circle is drawn for each of the models. The overlap between the circles represents overlap between the models. The numbers represent the number of unique PBs in a subset. For example, in **Figure 4b,** 28 PBs are only present in Replicate 1, whereas 8 PBs are shared between Replicate 2 and Replicate 3, and 108 PBs are present in all three replicates.

#### 9.2. Plotting distances

Clicking 'Plot distances' plots the distances of the PBs in the available set in a box plot combined with a strip plot (**Figure 4c**). Data are grouped per PB model. The plot can be saved by clicking the save icon 🖫 in the plot window.

#### 9.3. Finding shortest PBs

##### 9.3.1. Rationale

When one peptide pair yields multiple PBs, in the case of homomultimeric complexes like fibrin.pdb in **Figure 3,** for instance, it can be useful to take only the shortest PBs per peptide pair into account, since these are often more likely to be correct. However, it is important to realize that this procedure introduces bias.

##### 9.3.2. Procedure

Upon clicking 'Find shortest', the distance by which the PBs are allowed to differ can be specified (**Figure 4d**). For instance, when the specified distance is 1 Å, and the shortest PB of a peptide pair is 5 Å, PBs with a distance of at most (5 + 1 =) 6 Å are also included in the shortest PBs for that peptide pair. After clicking 'OK', a new PB model is generated, called *chosen_name*_shortest.pb, where *chosen_name* represents the name under 'Chosen name'. For instance, the PB model demo_1_shortest.pb in **Figure 2** resulted from applying the 'Find shortest' function to demo_1.pb.

#### 9.4. Customizing names

By default, the labels in plots, as well as the names of models generated via the Analyze window are equal to the name(s) of the model(s) in the available set. However, the name(s) to be used can be changed in the 'Customize name' panel; names under 'Chosen name' can be edited after double clicking them (**Figure 4a**).

#### 9.5. Updating distances

Clicking 'Update distances' updates the distances in the mapping information file. This is useful when PBs connecting two models are present, and the model's positions have been moved with respect to one another, thus changing the distances of inter-model PBs. If none of the selected PBs are represented in a mapping information file, as is the case when a PB's model was not created with XMAS, this button is disabled.

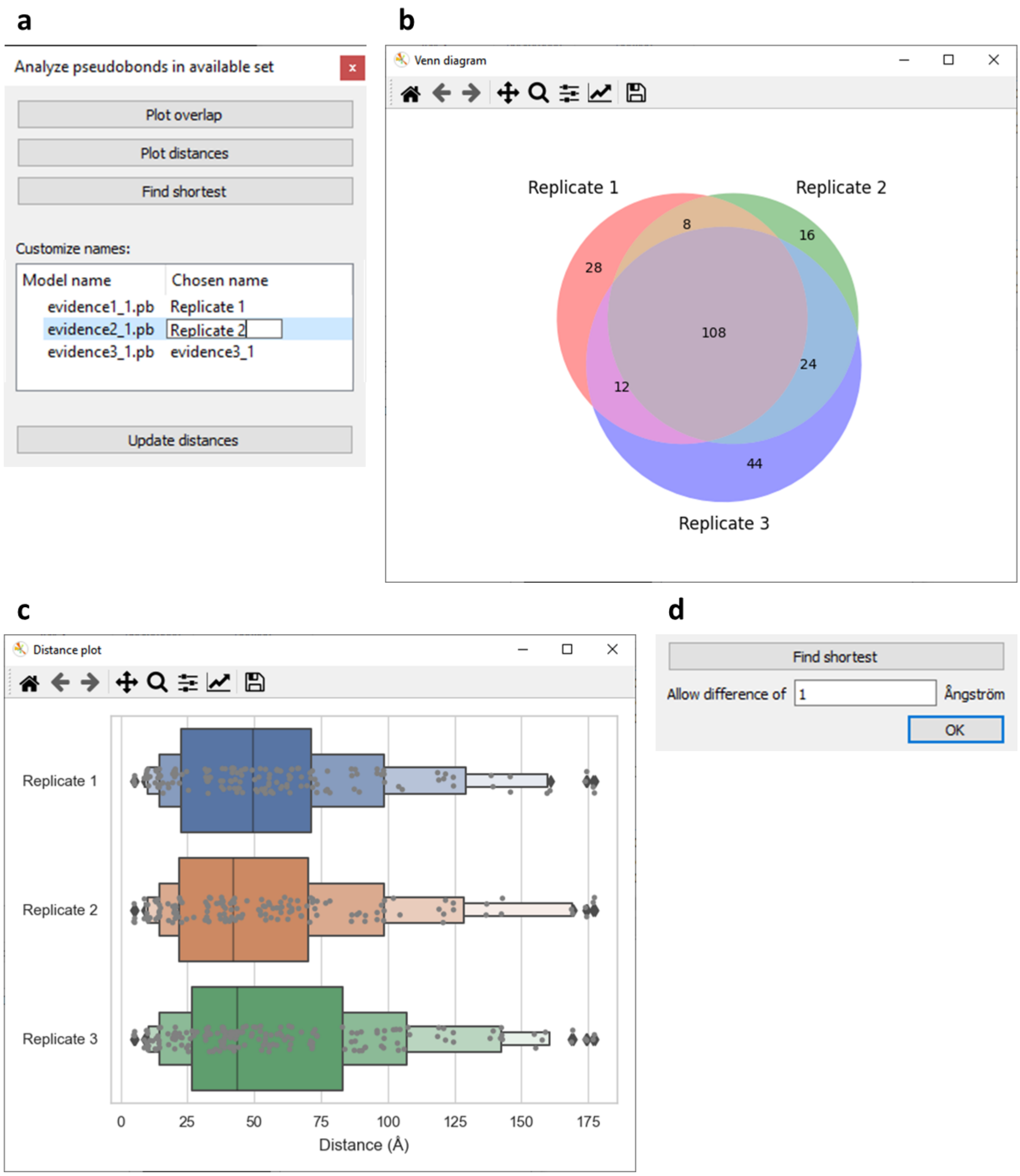

**Figure 4. Functionalities of the Analyze window.**

**a.** Analyze window opened when PB models evidence1_1.pb, evidence1_1.pb, and evidence1_1.pb were selected. **b.** Venn diagram created by clicking 'Plot overlap' in the Analyze window. **c.** Distance plot created by clicking 'Plot distances' in the Analyze window. **d.** Additional content appearing in the Analyze window upon clicking 'Find shortest', which enables specifying the allowed difference in distance between the shortest PB from a peptide pair and other PBs from that same peptide pair. PBs within the maximum allowed distance are included in the generated PB model.

#### 10. Exporting crosslinks

The 'Export' button opens the Export window, which contains components that enable specifying a subset to export, and the file format(s) of the thereby generated files (**Figure 5a**). The subset is specified by excluding specific PBs from the available set. Clicking 'Export subset' initializes the export.

#### 10.1. Specifying subset

Criteria that can be used to in- or exclude PBs from the exported subset are the molecular model(s) that they belong to, their crosslink type, distance, confidence score, and quantity.

##### 10.1.1. Molecular models

The 'Molecular models' panel contains all molecular models that the PBs in the available set belong to. For example, the PBs in the available set from **Figure 5a** are all between atoms from fibrin.pdb, so fibrin.pdb is the only model present in the panel. PBs that (partially) belong to a model that is unchecked in this panel are excluded from the exported subset.

##### 10.1.2. Crosslink type

In the 'Links' panel, three types of links are specified. A PB always belongs to one of these link types, depending on the atoms that it connects:

- In **intralinks**, connected atoms belonging to the same chain.
- In **chain interlinks**, connected atoms belonging to different chains in the same model.
- In **model interlinks**, connected atoms belonging to different models.

PBs of a link type with an unchecked checkbox are excluded from the exported subset.

A link type is disabled in the 'Links' panel when it conflicts with the models in the 'Molecular models' panel. For example, 'Model interlinks' is disabled in **Figure 5a**, because only one model is present in 'Molecular models', which means that there will be no PBs that connect atoms between different models. Likewise, if none of the molecular models contain multiple chains, 'Chain intralinks' will be disabled.

##### 10.1.3. Distance

PBs can be excluded from the exported subset based on their distance in Å, using a cut-off slider for distance. When a PB length violates the distance slider's allowed range, it is not displayed, allowing interactive visualization upon manipulation of the slider.

##### 10.1.4. Confidence score

To exclude PBs from the exported subset based on their peptide pair's confidence score in the evidence file, a cut-off slider for this score is used. If one or multiple PBs in the available set are not linked to a score, the score slider is disabled. A PB is only linked to a score if it was created by mapping an evidence file with XMAS. Consequently, opening a new .pb file with ChimeraX itself results in PBs without scores and a disabled score slider.

##### 10.1.5. Quantity

Two dropdown menus below 'Quantity' enable defining how many times a PB needs to be present in the available set in order to be included in the exported subset. A choice is made between 'least' and 'most' in the left dropdown menu, and the number of PB models is chosen in the right dropdown menu. For example, if only the PBs that are present in three out of three PB models ought to be included, 'least' should be selected in the left dropdown menu, and '3' in the right (**Figure 5a**). In this manner, only the PBs that are present in at 'least 3 out of 3 PB models' are included. Conversely, selecting 'most' and '1' results in including only PBs that are present at 'most 1 out of 3 PB models', i.e. the PBs that are uniquely present.

#### 10.2. Specifying file format

The subset can be saved as a PB file ('.pb' checked), a DisVis restraints file ('DisVis' checked), or both. Upon clicking 'Export subsets', a window will appear that enables choosing the storage location(s) and name(s) of the created file(s).

##### 10.2.1. PB file

When '.pb' is checked, a new PB file will be created, containing the chosen subset of PBs. These PBs will also be opened as a new PB model in the ChimeraX session.

##### 10.2.2. DisVis restraints file

When 'DisVis' is checked, a new restraints file will be created, containing the chosen subset of PBs. This file can be used as restraints file in a DisVis job. Restraints files differ from PB files, because the atom specifiers required for DisVis are different than those required for ChimeraX. Moreover, whereas the order of the atom specifiers is irrelevant in a PB file, DisVis requires all first atoms to be from one model (called 'fixed chain' in DisVis), and all second atoms from another model (called 'scanning chain'). Lastly, a minimum and maximum distance has to be specified for the DisVis restraints file. Before the file can be saved, a DisVis window appears (**Figure 5b**) in which the fixed and scanning chains should be specified, and the minimum and maximum distances can be assigned. See 12. Integrative modeling for more information on integrating XMAS and DisVis.

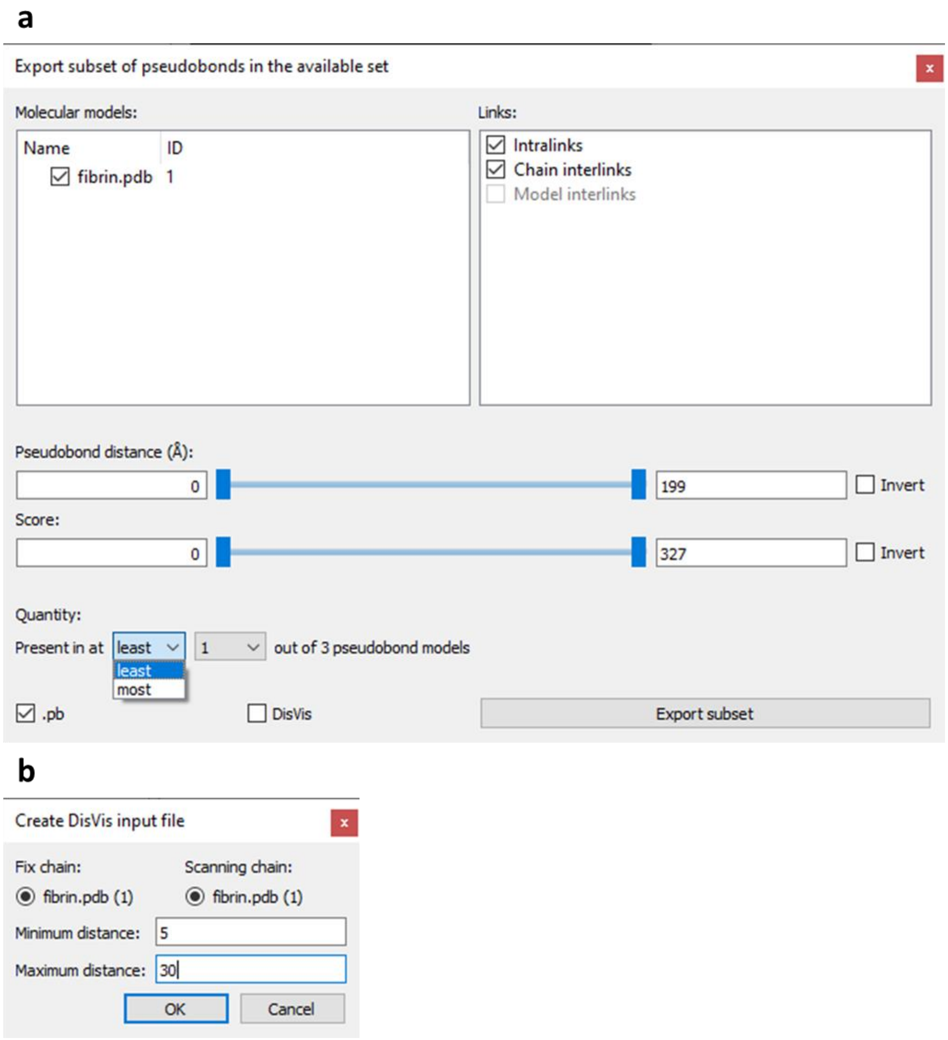

**Figure 5. Functionalities of the Export window.**
**a.** Example of an Export window. **b.** Example of a DisVis window.

### 11. Visualizing crosslinks

The ‘Visualize’ button opens a window with several options to change the appearance of the available set of PBs. The functions of the Visualize window can be subdivided in two categories: value-based coloring and customized styling (**Figure 6a**). The changes made can be discarded by clicking ‘Cancel’, or accepted by clicking ‘Apply’.

#### 11.1. Value-based coloring

This functionality entails coloring PBs according to their distances and confidence scores. For both types of values, PBs can be colored according to a gradient and a cut-off range. Color gradients are mutually exclusive, so when a gradient checkbox is checked, the other gradient checkbox is automatically unchecked. However, one color gradient can be used simultaneously with both cut-off options.

##### 11.1.1. Gradient coloring

Upon checking a gradient checkbox, the PBs are colored according to a color gradient that corresponds to the range of values. The colors and their corresponding values are displayed in a Color Key model at the bottom right corner of the ChimeraX model display area. The lowest value is set to 0 for the distance and confidence score gradient. The maximum distance gradient value is set to the maximum value found in the available set. For the confidence score, on the other hand, the maximum gradient value is set to 200, so that all PBs with a score of 200 or higher get the same color.

##### 11.1.2. Cut-off coloring

Upon checking a cut-off checkbox, a range can be specified using cut-off sliders for distance and confidence score. PBs with a distance or score that violates the range specified with the slider are colored differently than the other PBs. The colors used are described in 11.2. Customized styling.

#### 11.2. Customized styling

This functionality entails customizing color, radius, and number of dashes of the PBs. Colors and radii for PBs that are within the cut-off ranges of 11.1.2. Cut-off coloring (referred to as ‘Main’) can be set differently than those outside of these ranges (referred to as ‘Cut-off’). The number of dashes can only be applied to complete PB models. Consequently, the number of dashes change for all models that have at least one PB in the available set. By default, radii are 0.5 and the number of dashes is 8.

#### 11.3. Examples

Examples of images that can easily be rendered using the Visualize dialog are shown in **Figure 6b**.

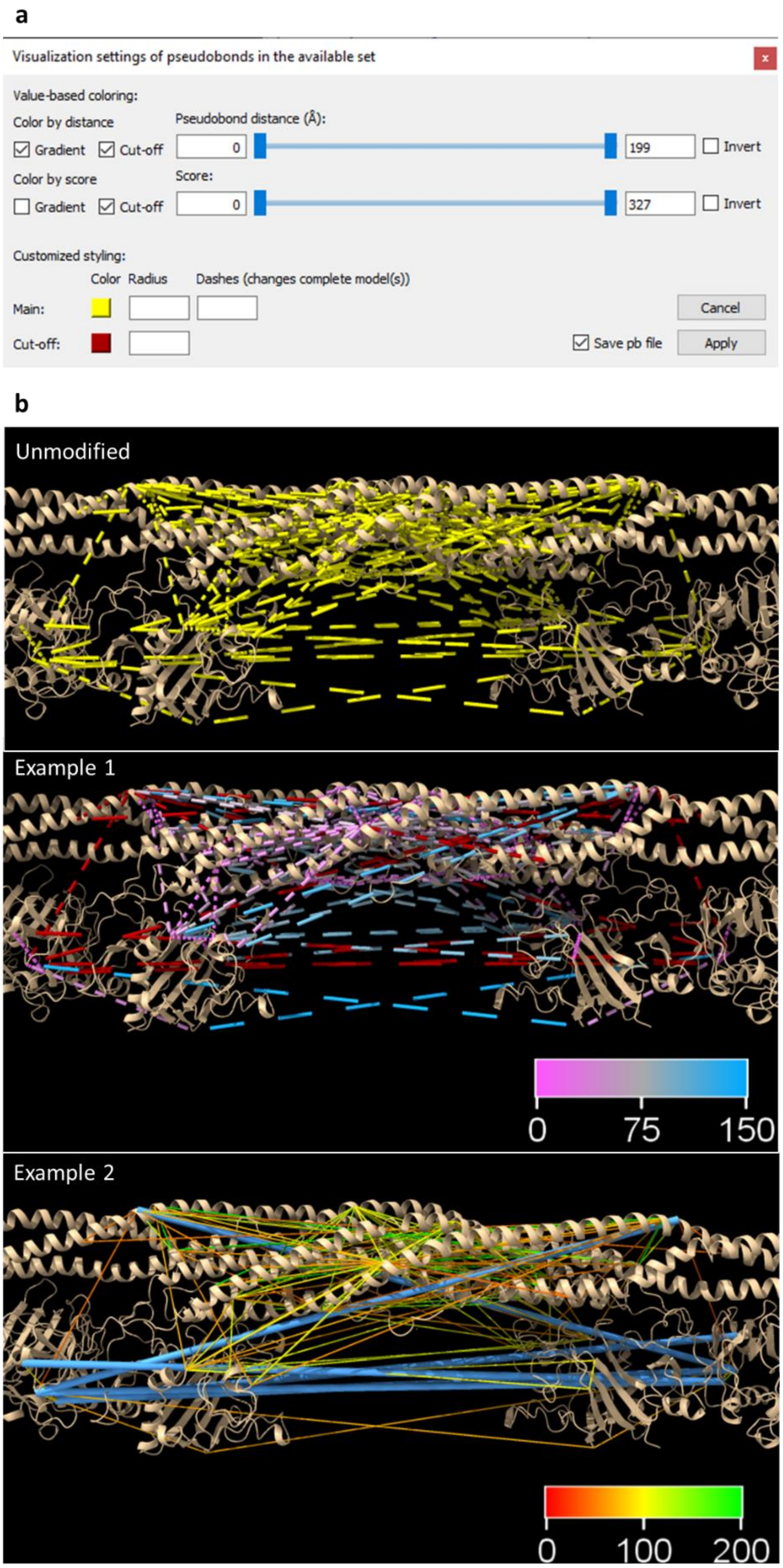

**Figure 6. Functionalities of the Visualize window.**

**a.** Example of a Visualize window. **b.** Examples of images created with the aid of the Visualize window. Example 1 was created by applying a distance gradient and a cut-off color (red) for distance and score cut-offs. The color key model represents the PB distances in Å and their corresponding color. Example 2 was created by applying a score gradient and the same cut-offs as in Example 1. The cut-off color was changed to blue. Moreover, the radii of 'Main' PBs were decreased, whereas the radii of 'Cut-off' PBs were increased. The number of dashes was set to 1.

### 12. Integrative modeling

XMAS supports integration with structural modeling platform DisVis. NB what is called a model in ChimeraX, is called a chain in DisVis. The term model is henceforth used in this section.

The restraints file containing the crosslinks in the DisVis-required format can be generated with the XMAS Export window. Additionally, the output of a DisVis job can be processed with the Integrate window (**Figure 7a**), which is opened by clicking 'Integrate' in the XMAS main window. The Integrate window enables DisVis output folder selection, reading and opening the relevant contents of this folder in ChimeraX, interactive visualization of restraint distances and z-scores, and z-score based restraints selection.

#### 12.1. Selecting an output folder

To select a DisVis output folder, the path to this folder is inserted in the text box, or 'Select folder' is clicked. The folder should always contain (**Figure 7b**):

- The **fixed and scanning models** as Protein Data Bank (PDB) files
- The **restraints** file that was used as input
- The DisVis **log** file; disvis.txt
- The **accessible interaction space** file; accessible_interaction_space.mrc
- The **z-scores** file; z-score.out

#### 12.2. Reading and opening relevant content

When the DisVis output folder is selected, and 'Use folder' is then clicked, XMAS will open the fixed and scanning models from their PDB files in the ChimeraX session. Additionally, each restraint in the restraints file is visualized as a PB. The specified minimum and maximum distances are also read from the restraints file. Moreover, the accessible interaction space is opened from accessible_interaction_space.mrc. Next, the z-scores are read from z-scores.out, and each z-score is linked to the PB that reflects the corresponding distance restraint. A cut-off slider for the z-scores appears in the Integrate window. Unlike other XMAS cut-off sliders, the initial minimum value is equal to the lowest z-score present, instead of zero.

#### 12.3. Interactive visualization of restraint parameters

After the DisVis output folder has been read, two relevant parameters, namely distances and z-scores, can be inspected visually.

##### 12.3.1. Distances

The PBs from the restraints file are colored either yellow or red, depending on whether they violate the specified minimum or maximum distance. Distances of yellow PBs are within the specified range, whereas those of red PBs are outside the range. The coloring will be adjusted automatically if models change position with respect to each other.

##### 12.3.2. Z-scores

When manipulating the z-score slider, PBs with a z-score outside the chosen range will be hidden from view.

#### 12.4. Z-score based restraints selection

To select the restraints within a chosen z-score range, the z-score slider is set to this range, and 'OK' is clicked. XMAS then writes a PB file containing only the PBs with an appropriate z-score. The written file is stored in a new folder, Selected_restraints, which is created inside the DisVis output folder used.

The file is named restraints_selected.pb, where *restraints* represents the name of the DisVis restraints file without its file extension ('.txt').

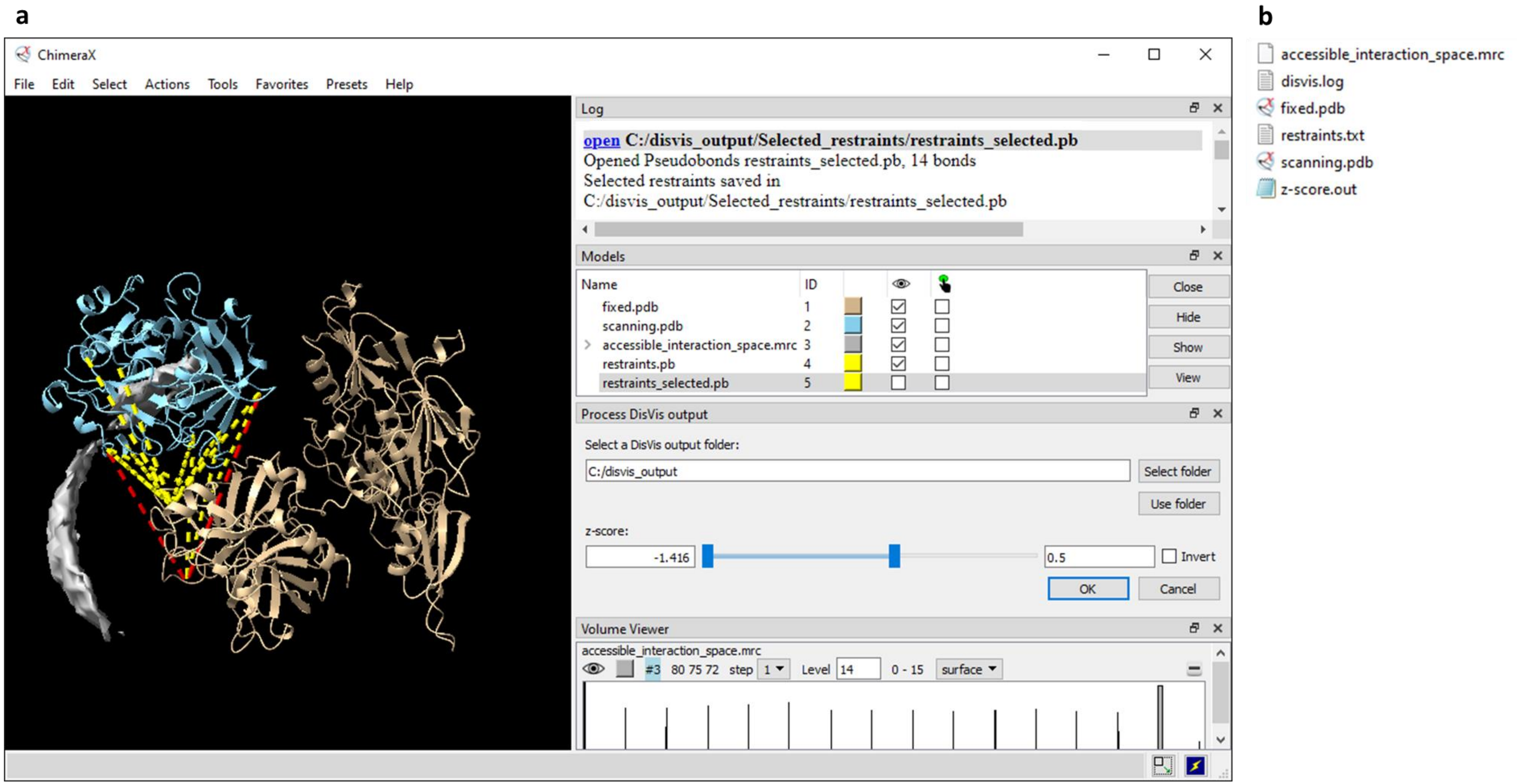

**Figure 7. Processing DisVis output.**

**a.** ChimeraX session in which the XMAS Integrate window (titled 'Process DisVis output') is used. After selecting the DisVis output folder (C:/disvis_output), 'Use folder' was clicked by the user. This prompted the opening of fixed.pdb, scanning.pdb, and accessible_interaction_space.mrc from the folder, and the appearance of the ChimeraX 'Volume Viewer' window, in which the user set the level at 14 consistent restraints. XMAS also created the PB model restraints.pb in this process. After the user specified a maximum z-score of 0.5 with the slider, and clicked 'OK', XMAS generated the PB file and model restraints_selected.pb, which contained only the PBs with a z-score of 0.5 or lower. In the log, the storage location of the generated PB file is mentioned. NB restraints_selected.pb is hidden from view in the session, the only visible PB model being restraints.pb. **b.** Files in the DisVis output folder used in **a**.

### 13. Recurring principles

Two principles occurring in multiple components of XMAS are the available set of PBs, cut-off slider, and DisVis job.

#### 13.1. Available set

The principle of the available set applies to the Analyze, Export, and Visualize windows. This set comprises the PBs that were selected at the moment that a window was opened. Operations performed in the window will exclusively influence the PBs in the available set, unless otherwise stated. NB changing the selection when an Analyze, Export, or Visualize window is already open, will not change the available set of PBs. Hence, a new window needs to be opened to use a different set. Furthermore, non-selected PBs in a partially selected model are excluded from the available set.

#### 13.2. Cut-off slider

Cut-off sliders are found in the Export, Visualize, and Integrate windows. These sliders dictate the range of values that a PB is allowed to have. The value covered by a slider can be distance, confidence score, or z-score. The minimum and maximum cut-off values are visible in the text boxes on the left and right side of the slider, respectively. Initially, the minimum value is zero, and the maximum value is equal to the highest value found in the set of PBs, unless otherwise stated. The cut-off values can be adjusted by typing the required minimum and maximum values in their text boxes. Moreover, they can be adjusted by moving the left and right slider handles, or the bar between those, using the cursor. By default, all values in the range between and including the cut-off values are considered allowed. The 'Invert' checkbox is checked to change the allowed range, to values equal to or lower than the minimum cut-off value, and equal to or higher than the maximum cut-off value. When multiple cut-off sliders are present in the same window, the allowed ranges of all sliders are applied.

#### 13.3. DisVis job

The input for a DisVis job comprises two models and crosslinks between these model. One model is defined as 'fixed chain' and the other as 'scanning chain'. The crosslinks are referred to 'restraints' and are loaded into DisVis in a restraints file, which can be generated with XMAS. A vast number of scanning chain positions (relative to the fixed chain), in which the two structures can theoretically interact chemically, is sampled. For each sampled position, it is determined whether a restraint is consistent with the specified distance. The DisVis-generated output files that are relevant for XMAS are the accessible interaction space and z-score files. For further technical details, please read the DisVis manual.
